## Supplementary Figures for "Global coral genomic vulnerability explains recent reef losses"

**This PDF file includes:**

Text: Extended Methods for Mutations Area Relationship
Tables S1 to S6
Figs. S1 to S22

### Table of content

**Extended methods for mutations area relationship** **3**
**References** **4**
**Supplementary Tables** **5**
Table S1. Genomic datasets. 5
Table S2. Coefficients of models explaining genetic distances between samples. 6
Table S3. Overlapping adaptive signals. 7
Table S4. Gene ontology enrichment analysis. 12
Table S5. Coefficients of Acropora cover change models. 13
Table S6. Parameters of Mutations-Area Relationship Linear Mixed Model. 14
**Supplementary Figures** **15**
Figure S1. Workflow of the study. 15
Figure S2. Sequencing read mapping stats. 16
Figure S3. Distribution of SNPs across genomes. 17
Figure S4. Principal component analysis of sequencing and alignment statistics. 18
Figure S5. Principal component analysis of genotype matrices. 19
Figure S6. Shared SNPs and % of non-identical genotypes across datasets. 20
Figure S7. Genetic proximity of all samples from the Acropora datasets. 21
Figure S8. Heat stress variables distribution across datasets. 22
Figure S9. Missing values and allele frequencies statistics. 23
Figure S10. Cross-entropy estimation of the number of ancestral populations. 24
Figure S11. Validation of overlapping adaptive signals on an independent Acropora dataset.25
Figure S12. Cross validation of the environmental genomic model. 26
Figure S13. Performance of models explaining Acropora cover decline across Indo-Pacific. 27
Figure S14. Worldwide distribution of Acorpora records from the Ocean Biodiversity Information System (OBIS). 28
Figure S15. Measured and projected maximal annual Degree Heating Week. 29
Figure S16. Temporal projections of heat exposure, frequency of heat-adapted genotypes and genomic vulnerability. 30
Figure S17. Acropora mutations-area relationships (MAR). 31
Figure S18. Acropora z-mar coefficients. 32
Figure S19. Theoretical Allele Frequency Spectrum for a panmictic population. 33
Figure S20. Z-mar coefficient variation by limiting the number of sampling sites. 34
Figure S21. Genetic extinction by area loss simulations. 35
Figure S22. Heat-adapted reefs in conservation and management strategies in 2020. 36

### Extended methods for mutations area relationship

We used the Mutations-Area Relationship (MAR) (Exposito-Alonso et al. 2022) framework to characterize how the genetic diversity of an *Acropora* population scales with the area occupied by the population. Using the same methods described in (Exposito-Alonso et al. 2022), we calculated the MAR for each *Acropora* dataset. In short, the raw genotype matrix was preliminarily filtered for missing observations - with a 20% threshold by individual and by SNP (Table S1D) - but not for allele frequencies. Focusing on 100 randomly picked SNPs from the genotype matrix, we repeated 50 times a subsampling procedure that consisted in drawing a square of random size across the spatial range of the dataset, retrieving the sampling sites falling within this square region, and calculating the number of mutations across the colonies from such sampling sites. The subsampling procedure was then replicated with ten different subsets of SNPs, yielding at total of 500 subsamples per dataset (Figure S9).

We then built a Bayesian linear mixed model to characterize the scaling between area and mutations across the subsamples of all the *Acropora* datasets. The model was built using the R-package MCMCglmm (v. 2.30) (Hadfield 2010), with log-transformed number of mutations as response variable, and log-transformed square area as main explanatory variable. Additional co-variables - such as identifiers of datasets and genomic subsamples, spatial autocorrelation indices, and mean sequencing depth - were included in the model after a stepwise construction to optimize the goodness of fit (according to the deviance information criterion - DIC; Table S5). This final model returned the z-mar parameter - *i.e.,* the coefficient of regression between log-transformed area and number of mutations - which summarizes the strength of the MAR. The z-mar estimated for *Acropora* was 0.14 [IC=0.07-0.19] (Figure S10). Two datasets showed a z-mar significantly lower or higher than the overall *Acropora* estimate: *A. tenuis* from the GBR (z-mar=0.06 [IC=0.00-0.11]) and *A. millepora* from New Caledonia (z.mar=0.23 [IC=0.16-0.28]), respectively.

As a comparison, we also calculated the z-mar expectation in each dataset under a panmictic population scenario. This was done by calculating MAR using a simulated genotype matrix, where individual genotypes were sampled from an allele frequency spectrum expected under panmixia (Figure S11). The simulated panmictic $z_{MAR}$ were systematically lower than the real z-mar (Figure S10)

The average *Acropora* z-mar was substantially smaller compared to the only z-mar available for corals to date, estimated for an *A. millepora* population from the GBR (z-mar=0.24 [0.20-0.28]) (Exposito-Alonso et al. 2022). We hypothesized that these low z-mar coefficients might be related to the relatively low number of sampling sites (<10) used in the majority of the *Acropora* datasets we analyzed. To further investigate this potential bias, we repeated the MAR subsampling procedure while limiting the maximal number of sampling sites, and observed that indeed increasing the number of sampling sites progressively raised the z-mar until a saturation point (Figure E8). Using the nlme R-package (v. 3.1) (Pinheiro et al. 2013), we designed a non-linear mixed model to characterize the increase of *Acropora* z-mar as a function of the number of sampling sites (while controlling for dataset effect as a random factor), and estimated an overall z-mar saturation point of 0.31 [IC=0.20-0.42].

We then asked whether the saturated *Acropora* z-mar parameter can predict genetic loss by different scenarios of habitat loss in the studied populations (Figure E9). For that, we computed stochastic simulations of extinction, where sampling locations were progressively removed from every dataset. As the sampling locations were removed, we kept track of the decrease in the number of mutations. We tested different spatial patterns of extinction: random (extinction randomly hits sampling sites; Figure E9A), radial (extinction starts at a random site, then propagates to the neighboring sites; Figure E9B), equatorial (extinction propagates from low to high latitudes; Figure E9C) and polar (extinction propagates from high to low latitudes, Figure E9D). In all the datasets, the four extinction scenarios aligned with the rates of genetic loss predicted by the saturated *Acropora* z-mar.

### Supplementary Tables

#### Table S1. Genomic datasets.

Shown are datasets-specific parameters for different processing steps of the study: (A) Dataset characteristics, (B) Filtering thresholds and outputs of reads quality check and mapping, (C) Parameters and outputs of the genotype-by-environment association analysis, (D) output of the mutations area relationship analysis.

| Bioproject ID | PRJNA593014 | PRJNA702071 | PRJNA665778 | PRJDB4188 | PRJEB37470 | PRJNA434194 | PRJNA750412 |
|---|---|---|---|---|---|---|---|
| **Authors** | **Fuller et al., 2020** | **Selmoni et al., 2021** | **Drury et al., 2017** | **Shinzato et al., 2015** | **Cooke et al., 2020** | **Matz et al., 2018** | **Torquato et al., 2022** |
| **A) Datasets characteristics** | | | | | | | |
| **Species** | *Acropora millepora* | *Acropora millepora* | *Acropora cervicornis* | *Acropora digitifera* | *Acropora tenuis* | *Acropora millepora* | *Acropora downingi* |
| **# unique coral genotypes** | 240 | 187 | 179 | 155 | 150 | 85* | 61 |
| **# sampling locations** | 12 | 20 | 22** | 12 | 5 | 5 | 10 |
| **Samples per site** | 10 to 20 | 10 to 15 | 1 to 20 | 10 to 20 | 30 | 20-30 | 1 to 10 |
| **Region** | Central GBR | New Caledonia | Florida Reef Tract | Ryukyu Archipelago | Central GBR | GBR | Persian Gulf |
| **Spatial range** | 200 km | 400 km | 400 km | 1500 km | 500 km | 1000 km | 800 km |
| **Sequencing strategy** | WGS (low-coverage) | RAD-Seq (DArT-seq) | RAD-Seq (GBS) | WGS | WGS (low-coverage) | RAD-Seq (2bRAD) | RAD-Seq |
| **Sequencing machine** | Illumina HiSeq 4000 | Illumina HiSeq 2500 | Illumina HiSeq 2500 | Illumina HiSeq 2000 | Illumina HiSeq 2500 | Illumina HiSeq 2000 | Illumina HiSeq 2000 |
| **B) Filtering and SNP calling** | | | | | | | |
| **Hard clipping (5' - 3')** | 11 - 5 | 15 - 3 | 2 - 2 | 5 - 2 | 15 - 3 | 1 - 2 | 6 - 2 |
| **# of corals after quality filtering** | 234 | 167 | 128 | 154 | 139 | 85 | 56 |
| **# of SNPs** | 16,338 | 42,864 | 9,303 | 99,810 (23,854,772 before subsampling) | 98,285 (1,375,995 before subsampling) | 64,947 | 32,537 |
| **# of coral after outlier filtering** | - | 163 | 76 | 140 | 117 | 79 | 38 |
| **# sampling sites after filtering** | - | 18 | 14 | 9 | 4 | 5 | 8 |
| **Pairwise nucleotide diversity (π/bp)** | | 0.21% | 0.45% | 0.59% | 0.91% | 0.14% | 0.33% |
| **C) Genotype-environment association analysis** | | | | | | | |
| **# of SNPs retained after all. Fr. filtering** | - | 9,385 | 2,773 | 15,537 | 15,855 | 25,597 | 17,392 |
| **# of colonies retained after allele fr. filtering** | - | 156 | 69 | 140 | 116 | 79 | 35 |
| **# of latent factors in LFMM** | - | 4 | 4 | 1 | 1 | 1 | 1 |
| **LFMM genomic inflation factor (λ) before calibration** | - | 1.13 | 1.22 | 1.19 | 0.99 | 1.29 | 1.39 |
| **LFMM genomic inflation factor (λ) after calibration** | | 1 | 1 | 1 | 1 | 1 | 1 |
| **# of overlapping genomic windows (q<0.1)** | - | 39 | 18 | 65 | 56 | 68 | 42 |
| **D) Mutations Area Relationship** | | | | | | | |
| **# of SNPs retained after filtering** | - | 40,441 | 8,474 | 85,420 | 68,953 | 64,430 | 32,441 |
| **# of colonies retained after filtering** | - | 155 | 63 | 133 | 90 | 79 | 34 |
| **Raw z-mar** | - | 0.23 [0.16-0.28] | 0.13 [0.07-0.19] | 0.16 [0.10-0.21] | 0.06 [0.00-0.11] | 0.10 [0.04-0.16] | 0.15 [0.09-0.21] |
| **Panmictic z-mar** | - | 0.07 [0.05-0.10] | 0.10 [0.07-0.13] | 0.05 [0.02-0.07] | 0.02 [0.00-0.04] | 0.05 [0.03-0.08] | 0.08 [0.05-0.11] |

* includes samples collected in 2002, excluding 24 samples collected in 2009 and 2014.

** samples collected within 5km of distance were assigned to the same sampling site.

| Response variable | PCoA-1 | | | PCoA-2 | | |
|---|---|---|---|---|---|---|
| | *P (Longitude)* | *P (Latitude)* | *AIC* | *P (Longitude)* | *P (Latitude)* | *AIC* |
| **Full model** | 2.02E-22 | 1.26E-05 | -1349.13 | 0.733068 | 0.024438 | -1290.71 |
| **Model without Species** | 2.02E-22 | 1.26E-05 | -1351.13 | 0.960467 | 0.00847 | -1290.09 |
| **Model without Dataset** | 1.64E-13 | 4.65E-01 | -1337.97 | 0.170672 | 0.126376 | -1290.21 |
| **Model with Lon/Lat** | *NA* | *NA* | -1342.43 | *NA* | *NA* | -1290.91 |

**Table S2. Coefficients of models explaining genetic distances between samples.**

Shown are the coefficients of linear mixed models used to explain the genetic principal coordinate axes (PCoA-1 and PCoA-2) of the six *Acropora* datasets. For every model, shown are the p-value associated with the effects of longitude and latitude, and the overall Akaike Information Criterion (AIC). The full model separates samples based on geography (longitude and latitude), species and dataset. The alternative models remove each one of these factors.

**Table S3. Overlapping adaptive signals.**

Shown is the list of the 85 genomic windows where genotype-environment associations were repeatedly found in different datasets. For every genomic window, the table displays the chromosome or the contig on the *A. millepora* reference genome, the start and the end of the window, q-value of the overlap analysis from PicMin (*23*), the datasets were overlapping single nucleotide polymorphisms (SNPs) were detected and the position of the most significant SNP by dataset. The table also displays information on genes located in the genomic window: number of genes (#), gene identifier (ID), gene start and end, and annotated proteins. The table also summarizes the results of the differential gene expression analysis of genes (DGE) (*34*), where "+" indicates up-regulation of the gene under heat exposure, "-" indicates down-regulation, "ns" a non-significant change, and "?" an unknown change.

| | Genomic windows | | | SNPs | | Genes | | | | | |
|---|---|---|---|---|---|---|---|---|---|---|---|
| | Chromosome/ contig | Start- End | q | Datasets | Position | # | ID | DGE | Start | End | Annotated Protein |
| 1 | chr1<br>NC_058066.1 | 10,290,001-<br>10,290,001 | 0.099 | Selmoni<br>Matz<br>Torquato | 10,292,969<br>10,292,819<br>10,293,302 | 1 | LOC114961598 | ns | 10,282,587 | 10,297,218 | Fibronectin |
| 2 | chr1<br>NC_058066.1 | 20,040,001-<br>20,040,001 | 0.055 | Selmoni<br>Shinzato<br>Cooke<br>Torquato | 20,045,440<br>20,044,813<br>20,044,377<br>20,046,394 | 1 | LOC114956149 | ns | 20,031,891 | 20,059,549 | Protein mesh |
| 3 | chr1<br>NC_058066.1 | 22,770,001-<br>22,770,001 | 0.068 | Cooke<br>Matz<br>Shinzato<br>Torquato | 22,772,396<br>22,778,603<br>22,776,094<br>22,776,461 | 1 | LOC114967796 | ? | 22,770,690 | 22,772,471 | PiggyBac transposable element-derived protein 4 |
| 4 | chr1<br>NC_058066.1 | 28,010,001-<br>28,010,001 | 0.08 | Matz<br>Shinzato<br>Cooke | 28,017,048<br>28,017,768<br>28,016,099 | 1 | LOC114950224 | ns | 28,017,132 | 28,019,790 | - |
| 5 | chr1<br>NC_058066.1 | 36,370,001-<br>36,370,001 | 0.068 | Cooke<br>Selmoni<br>Matz<br>Shinzato | 36,375,293<br>36,379,850<br>36,378,841<br>36,378,433 | 1 | LOC114966583 | - | 36,304,385 | 36,381,911 | Adhesion G-protein coupled receptor V1 |
| 6 | chr1<br>NC_058066.1 | 37,010,001-<br>37,010,001 | 0.068 | Cooke<br>Shinzato<br>Selmoni<br>Matz | 37,011,109<br>37,011,622<br>37,018,656<br>37,010,163 | 2 | LOC114960048<br>LOC114960103 | ?<br>ns | 37,011,445<br>37,004,037 | 37,017,233<br>37,010,653 | Protein artichoke<br>Leucine-rich repeat-containing protein 57 |
| 7 | chr1<br>NC_058066.1 | 37,300,001-<br>37,300,001 | 0.068 | Matz<br>Cooke<br>Torquato<br>Shinzato | 37,306,398<br>37,309,268<br>37,301,717<br>37,303,683 | 1 | LOC114972168 | ns | 37,271,633 | 37,311,277 | - |
| 8 | chr1<br>NC_058066.1 | 37,540,001-<br>37,540,001 | 0.09 | Shinzato<br>Matz<br>Selmoni<br>Cooke | 37,541,563<br>37,543,164<br>37,542,950<br>37,544,622 | 2 | LOC114953470<br>LOC114953439 | ns<br>? | 37,540,530<br>37,548,234 | 37,545,410<br>37,550,680 | -<br>Protein lin-41 |
| 9 | chr2<br>NC_058067.1 | 1,260,001-<br>1,260,001 | 0.077 | Selmoni<br>Shinzato<br>Matz | 1,260,718<br>1,268,457<br>1,268,425 | 2 | LOC114961811<br>LOC114962376 | ns<br>ns | 1,269,377<br>1,247,455 | 1,272,079<br>1,265,456 | Serine/threonine-protein kinase pim-3<br>Calcium transport protein 1 |
| 10 | chr2<br>NC_058067.1 | 3,670,001-<br>3,670,001 | 0.032 | Matz<br>Torquato<br>Cooke | 3,675,988<br>3,677,185<br>3,674,210 | 1 | LOC114955787 | + | 3,659,929 | 3,679,326 | Adhesion G protein-coupled receptor E2 |
| 11 | chr2<br>NC_058067.1 | 3,700,001-<br>3,700,001 | 0.055 | Shinzato<br>Selmoni<br>Matz | 3,709,869<br>3,709,557<br>3,708,594 | 1 | LOC114955762 | + | 3,687,361 | 3,714,182 | Phosphatidylinositide phosphatase SAC2 |
| 12 | chr2<br>NC_058067.1 | 5,990,001-<br>5,990,001 | 0.068 | Shinzato<br>Torquato<br>Matz | 5,992,273<br>5,993,306<br>5,995,822 | 1 | LOC114972398 | + | 5,990,719 | 6,005,366 | Heat shock 70 kDa protein 9 |
| 13 | chr2<br>NC_058067.1 | 9,540,001-<br>9,540,001 | 0.055 | Drury<br>Matz<br>Selmoni | 9,540,633<br>9,540,606<br>9,544,095 | 2 | LOC114958799<br>LOC114958814 | ns<br>- | 9,548,089<br>9,514,067 | 9,573,614<br>9,540,383 | p53-induced death domain-containing protein 1<br>Kinesin-like protein KIF23 |
| 14 | chr2<br>NC_058067.1 | 16,800,001-<br>16,800,001 | 0.077 | Matz<br>Drury<br>Shinzato | 16,802,448<br>16,808,926<br>16,808,055 | 2 | LOC114976631<br>LOC114976630 | -<br>- | 16,804,607<br>16,798,098 | 16,809,847<br>16,802,764 | Nucleotidyltransferase MB21D2 |

| | Genomic windows | | | SNPs | | Genes | | | | | |
|---|---|---|---|---|---|---|---|---|---|---|---|
| | **Chromosome/ contig** | **Start- End** | **q** | **Datasets** | **Position** | **#** | **ID** | **DGE** | **Start** | **End** | **Annotated Protein** |
| | | | | Selmoni | 16,800,952 | | | | | | Nucleotidyltransferase MB21D2 |
| 15 | chr2<br>NC_058067.1 | 21,030,001-<br>21,030,001 | 0.096 | Matz<br>Selmoni<br>Torquato<br>Cooke | 21,031,328<br>21,031,398<br>21,031,387<br>21,035,858 | 2 | LOC114965334<br>LOC114965340 | ns<br>? | 21,031,136<br>21,026,209 | 21,046,403<br>21,031,081 | -<br>- |
| 16 | chr2<br>NC_058067.1 | 29,350,001-<br>29,350,001 | 0.096 | Selmoni<br>Matz<br>Shinzato | 29,356,508<br>29,351,800<br>29,353,020 | 2 | LOC114959064<br>LOC114959046 | -<br>- | 29,358,322<br>29,349,235 | 29,370,911<br>29,355,250 | Cyclic nucleotide-binding domain-containing protein 1<br>Tetratricopeptide repeat protein 28 |
| 17 | chr2<br>NC_058067.1 | 34,260,001-<br>34,260,001 | 0.032 | Matz<br>Cooke<br>Shinzato<br>Torquato<br>Selmoni | 34,267,757<br>34,264,302<br>34,260,981<br>34,265,179<br>34,265,299 | 1 | LOC114955874 | ns | 34,250,276 | 34,269,826 | Tetratricopeptide repeat protein 28 |
| 18 | chr3<br>NC_058068.1 | 8,920,001-<br>8,920,001 | 0.068 | Torquato<br>Matz<br>Selmoni<br>Shinzato | 8,925,701<br>8,923,136<br>8,925,701<br>8,925,172 | 1 | LOC114965837 | - | 8,909,636 | 8,938,454 | WASH complex subunit 5 |
| 19 | chr4<br>NC_058069.1 | 440,001-<br>440,001 | 0.069 | Matz<br>Cooke<br>Shinzato | 445,838<br>443,286<br>440,029 | 1 | LOC122947032 | ns | 439,881 | 445,916 | - |
| 20 | chr4<br>NC_058069.1 | 3,060,001-<br>3,060,001 | 0.068 | Matz<br>Cooke<br>Drury | 3,068,565<br>3,068,695<br>3,068,895 | 4 | LOC122958784<br>LOC122958782<br>LOC122958780<br>LOC114961461 | ?<br>?<br>?<br>ns | 3,064,798<br>3,065,666<br>3,067,061<br>3,057,442 | 3,074,174<br>3,067,756<br>3,071,926<br>3,121,257 | -<br>-<br>-<br>- |
| 21 | chr4<br>NC_058069.1 | 9,090,001-<br>9,090,001 | 0.055 | Torquato<br>Shinzato<br>Cooke<br>Matz<br>Drury | 9,099,279<br>9,099,342<br>9,099,069<br>9,099,948<br>9,099,806 | NA | NA | ? | NA | NA | NA |
| 22 | chr4<br>NC_058069.1 | 12,040,001-<br>12,040,001 | 0.09 | Matz<br>Shinzato<br>Cooke | 12,041,461<br>12,049,516<br>12,042,377 | NA | NA | ? | NA | NA | NA |
| 23 | chr4<br>NC_058069.1 | 13,430,001-<br>13,430,001 | 0.068 | Matz<br>Selmoni<br>Shinzato | 13,433,943<br>13,430,204<br>13,439,104 | 1 | LOC114955057 | ? | 13,428,967 | 13,441,371 | - |
| 24 | chr4<br>NC_058069.1 | 16,120,001-<br>16,120,001 | 0.077 | Shinzato<br>Torquato<br>Matz | 16,125,066<br>16,124,565<br>16,124,708 | 1 | LOC114948286 | ? | 16,123,785 | 16,127,292 | Retrovirus-related Pol polyprotein from transposon 412 |
| 25 | chr4<br>NC_058069.1 | 16,390,001-<br>16,390,001 | 0.099 | Cooke<br>Shinzato<br>Matz<br>Drury | 16,398,586<br>16,398,577<br>16,394,277<br>16,396,153 | 1 | LOC114948146 | ns | 16,399,214 | 16,403,418 | - |
| 26 | chr4<br>NC_058069.1 | 21,340,001-<br>21,340,001 | 0.077 | Shinzato<br>Matz<br>Torquato | 21,343,863<br>21,348,294<br>21,347,554 | 2 | LOC114954496<br>LOC114954494 | ns<br>ns | 21,342,547<br>21,292,365 | 21,345,558<br>21,342,456 | -<br>Hemicentin-1 |
| 27 | chr4<br>NC_058069.1 | 27,320,001-<br>27,320,001 | 0.055 | Cooke<br>Shinzato<br>Matz<br>Selmoni | 27,326,142<br>27,327,999<br>27,327,135<br>27,325,110 | 2 | LOC114962131<br>LOC114962111 | ?<br>ns | 27,322,992<br>27,329,038 | 27,328,231<br>27,333,823 | ALK tyrosine kinase receptor<br>Beta-2 adrenergic receptor |
| 28 | chr5<br>NC_058070.1 | 1,190,001-<br>1,190,001 | 0.055 | Selmoni<br>Shinzato<br>Matz | 1,194,560<br>1,191,632<br>1,190,599 | 2 | LOC114948182<br>LOC114948177 | ?<br>ns | 1,196,659<br>1,188,421 | 1,199,852<br>1,195,727 | -<br>- |
| 29 | chr5<br>NC_058070.1 | 9,000,001-<br>9,000,001 | 0.055 | Matz<br>Cooke<br>Shinzato | 9,001,969<br>9,004,851<br>9,005,746 | 1 | LOC114951588 | ? | 9,003,175 | 9,004,632 | - |
| 30 | chr5<br>NC_058070.1 | 11,100,001-<br>11,100,001 | 0.09 | Cooke<br>Torquato<br>Matz | 11,109,730<br>11,109,282<br>11,104,509 | 5 | LOC122959092<br>LOC122959093<br>LOC122959094<br>LOC114950235<br>LOC122959730 | ?<br>?<br>?<br>?<br>? | 11,102,071<br>11,103,159<br>11,108,705<br>11,109,987<br>11,099,417 | 11,104,763<br>11,104,550<br>11,110,011<br>11,111,440<br>11,104,763 | -<br>-<br>-<br>-<br>- |
| 31 | chr5<br>NC_058070.1 | 20,580,001-<br>20,580,001 | 0.096 | Selmoni<br>Torquato<br>Matz | 20,584,895<br>20,584,895<br>20,585,723 | 1 | LOC114955657 | ns | 20,550,717 | 20,610,432 | Transmembrane protein KIAA1109 |

| | Genomic windows | | | SNPs | | Genes | | | | | |
|---|---|---|---|---|---|---|---|---|---|---|---|
| | Chromosome/ contig | Start- End | q | Datasets | Position | # | ID | DGE | Start | End | Annotated Protein |
| 32 | chr5<br>NC_058070.1 | 21,150,001-<br>21,150,001 | 0.055 | Torquato<br>Cooke<br>Selmoni<br>Shinzato | 21,155,842<br>21,152,447<br>21,155,816<br>21,157,496 | 2 | LOC122959746<br>LOC114969872 | ?<br>- | 21,150,186<br>21,148,002 | 21,150,557<br>21,173,165 | -<br>Cytoplasmic tR- 2-thiolation protein 1 |
| 33 | chr6<br>NC_058071.1 | 1,670,001-<br>1,670,001 | 0.09 | Selmoni<br>Shinzato<br>Matz<br>Torquato | 1,672,635<br>1,674,718<br>1,674,024<br>1,672,522 | 1 | LOC114967515 | ns | 1,674,175 | 1,681,593 | - |
| 34 | chr6<br>NC_058071.1 | 5,580,001-<br>5,580,001 | 0.068 | Shinzato<br>Cooke<br>Matz | 5,589,419<br>5,586,669<br>5,589,241 | 1 | LOC114957539 | ? | 5,586,232 | 5,588,134 | Uncharacterized protein K02A2.6 |
| 35 | chr6<br>NC_058071.1 | 12,700,001-<br>12,700,001 | 0.055 | Selmoni<br>Shinzato<br>Cooke | 12,700,467<br>12,709,531<br>12,705,382 | 2 | LOC122960374<br>LOC114973614 | ns<br>ns | 12,706,748<br>12,696,343 | 12,715,968<br>12,703,580 | Olfactomedin-like protein 2A<br>Olfactomedin-like protein 2A |
| 36 | chr6<br>NC_058071.1 | 14,570,001-<br>14,570,001 | 0.041 | Torquato<br>Shinzato<br>Selmoni | 14,570,807<br>14,571,010<br>14,570,851 | NA | NA | ? | NA | NA | NA |
| 37 | chr7<br>NC_058072.1 | 2,720,001-<br>2,720,001 | 0.055 | Shinzato<br>Torquato<br>Matz | 2,725,178<br>2,722,514<br>2,725,837 | 1 | LOC114977834 | ? | 2,727,256 | 2,729,799 | - |
| 38 | chr7<br>NC_058072.1 | 7,030,001-<br>7,030,001 | 0.068 | Cooke<br>Matz<br>Shinzato | 7,034,550<br>7,033,558<br>7,033,024 | 1 | LOC122960927 | ? | 7,039,454 | 7,041,109 | - |
| 39 | chr8<br>NC_058073.1 | 9,910,001-<br>9,910,001 | 0.068 | Cooke<br>Matz<br>Drury | 9,914,843<br>9,919,385<br>9,910,115 | 2 | LOC114974575<br>LOC114974573 | ns<br>ns | 9,914,336<br>9,909,255 | 9,915,505<br>9,925,645 | -<br>- |
| 40 | chr8<br>NC_058073.1 | 13,180,001-<br>13,180,001 | 0.097 | Torquato<br>Matz<br>Shinzato | 13,182,663<br>13,181,144<br>13,184,594 | 1 | LOC114968694 | - | 13,167,639 | 13,186,522 | Ubiquitin-like modifier-activating enzyme 1 |
| 41 | chr9<br>NC_058074.1 | 3,590,001-<br>3,590,001 | 0.055 | Torquato<br>Cooke<br>Shinzato | 3,597,175<br>3,590,672<br>3,597,439 | 2 | LOC114970030<br>LOC114970008 | ns<br>ns | 3,594,241<br>3,579,976 | 3,598,810<br>3,608,703 | -<br>Polyunsaturated fatty acid 5-lipoxygenase |
| 42 | chr9<br>NC_058074.1 | 4,430,001-<br>4,430,001 | 0.087 | Torquato<br>Cooke<br>Drury<br>Selmoni | 4,434,503<br>4,434,345<br>4,434,480<br>4,433,072 | 2 | LOC114970072<br>LOC114970097 | +<br>+ | 4,427,847<br>4,421,354 | 4,437,446<br>4,450,739 | Uromodulin<br>60S ribosomal export protein NMD3 |
| 43 | chr9<br>NC_058074.1 | 5,130,001-<br>5,130,001 | 0.084 | Cooke<br>Matz<br>Shinzato | 5,130,345<br>5,130,066<br>5,130,530 | 1 | LOC114960495 | - | 5,129,505 | 5,148,460 | Protein unc-45 homolog B |
| 44 | chr9<br>NC_058074.1 | 13,920,001-<br>13,920,001 | 0.032 | Selmoni<br>Cooke<br>Matz | 13,928,615<br>13,921,057<br>13,928,074 | 1 | LOC114976214 | ns | 13,928,331 | 13,951,911 | Glutamate receptor 7 |
| 45 | chr9<br>NC_058074.1 | 16,750,001-<br>16,750,001 | 0.081 | Cooke<br>Drury<br>Torquato | 16,759,788<br>16,759,033<br>16,752,501 | 1 | LOC114951044 | ns | 16,743,398 | 16,762,292 | - |
| 46 | chr10<br>NC_058075.1 | 3,260,001-<br>3,260,001 | 0.05 | Shinzato<br>Cooke<br>Matz | 3,266,584<br>3,265,502<br>3,266,440 | 1 | LOC114953722 | + | 3,248,813 | 3,265,797 | Protein SSUH2 homolog |
| 47 | chr10<br>NC_058075.1 | 3,270,001-<br>3,270,001 | 0.063 | Shinzato<br>Matz<br>Cooke | 3,276,088<br>3,275,457<br>3,273,730 | 1 | LOC114953700 | ns | 3,279,372 | 3,282,142 | Late endosomal/lysosomal adaptor and MAPK and MTOR activator 5 |
| 48 | chr10<br>NC_058075.1 | 14,450,001-<br>14,450,001 | 0.055 | Matz<br>Shinzato<br>Drury | 14,452,192<br>14,454,569<br>14,456,349 | 3 | LOC114963342<br>LOC114963347<br>LOC114963351 | ?<br>-<br>ns | 14,451,276<br>14,459,869<br>14,442,434 | 14,456,429<br>14,461,923<br>14,451,192 | Tripartite motif-containing protein 45<br>-<br>Polysaccharide biosynthesis domain-containing protein 1 |
| 49 | chr10<br>NC_058075.1 | 14,550,001-<br>14,550,001 | 0.097 | Matz<br>Shinzato<br>Cooke<br>Drury | 14,554,297<br>14,554,855<br>14,554,897<br>14,557,157 | 3 | LOC114963223<br>LOC114963202<br>LOC114963212 | -<br>+<br>- | 14,552,807<br>14,559,607<br>14,539,768 | 14,559,514<br>14,594,938<br>14,552,829 | Frizzled-8<br>Enolase 4<br>Leucine-rich repeat-containing protein 27 |
| 50 | chr10<br>NC_058075.1 | 17,570,001-<br>17,570,001 | 0.09 | Cooke<br>Matz<br>Shinzato | 17,570,705<br>17,578,090<br>17,579,394 | 1 | LOC114947682 | - | 17,578,092 | 17,592,389 | Acylamino-acid-releasing enzyme |
| 51 | chr10<br>NC_058075.1 | 18,370,001-<br>18,370,001 | 0.077 | Shinzato<br>Drury | 18,378,044<br>18,372,559 | 1 | LOC114953413 | - | 18,356,789 | 18,384,393 | E3 ubiquitin-protein ligase RNF123 |

| | Genomic windows | | | SNPs | | Genes | | | | | |
|---|---|---|---|---|---|---|---|---|---|---|---|
| | Chromosome/ contig | Start- End | q | Datasets | Position | # | ID | DGE | Start | End | Annotated Protein |
| | | | | Matz | 18,370,350 | | | | | | |
| 52 | chr11<br>NC_058076.1 | 8,540,001-<br>8,540,001 | 0.032 | Cooke<br>Selmoni<br>Shinzato | 8,543,635<br>8,548,604<br>8,544,491 | 1 | LOC114975581 | - | 8,517,855 | 8,546,063 | WD repeat and HMG-box D--binding protein 1 |
| 53 | chr11<br>NC_058076.1 | 14,800,001-<br>14,800,001 | 0.068 | Cooke<br>Shinzato<br>Drury | 14,809,552<br>14,801,782<br>14,808,658 | 1 | LOC114957149 | ns | 14,798,346 | 14,806,253 | SET domain-containing protein 14 |
| 54 | chr11<br>NC_058076.1 | 16,790,001-<br>16,790,001 | 0.096 | Shinzato<br>Matz<br>Drury | 16,795,226<br>16,794,615<br>16,798,343 | 1 | LOC114965395 | ? | 16,789,422 | 16,790,592 | - |
| 55 | chr12<br>NC_058077.1 | 1,740,001-<br>1,740,001 | 0.091 | Cooke<br>Matz<br>Torquato<br>Shinzato | 1,749,896<br>1,742,711<br>1,741,493<br>1,740,808 | 1 | LOC114956891 | ns | 1,730,631 | 1,759,363 | Lactadherin |
| 56 | chr12<br>NC_058077.1 | 4,900,001-<br>4,900,001 | 0.068 | Selmoni<br>Shinzato<br>Matz | 4,900,951<br>4,908,255<br>4,900,573 | 2 | LOC114974291<br>LOC114974126 | +<br>? | 4,900,904<br>4,907,920 | 4,903,832<br>4,910,262 | -<br>- |
| 57 | chr12<br>NC_058077.1 | 11,270,001-<br>11,270,001 | 0.096 | Torquato<br>Matz<br>Shinzato<br>Cooke | 11,271,053<br>11,270,917<br>11,278,506<br>11,273,716 | 2 | LOC114968482<br>LOC114948467 | ?<br>ns | 11,276,547<br>11,269,275 | 11,278,954<br>11,274,326 | -<br>- |
| 58 | chr12<br>NC_058077.1 | 12,910,001-<br>12,910,001 | 0.096 | Cooke<br>Matz<br>Shinzato | 12,918,298<br>12,916,013<br>12,912,348 | 2 | LOC114976680<br>LOC114976677 | ?<br>? | 12,918,017<br>12,909,998 | 12,921,173<br>12,913,164 | -<br>- |
| 59 | chr12<br>NC_058077.1 | 18,520,001-<br>18,520,001 | 0.099 | Shinzato<br>Matz<br>Selmoni | 18,524,756<br>18,522,348<br>18,525,873 | 3 | LOC114958632<br>LOC122965083<br>LOC114958635 | +<br>?<br>+ | 18,522,692<br>18,529,546<br>18,518,400 | 18,539,550<br>18,534,505<br>18,523,134 | H(+)/Cl(-) exchange transporter 7<br>-<br>Cerebellar degeneration-related protein 2-like |
| 60 | chr13<br>NC_058078.1 | 10,880,001-<br>10,880,001 | 0.091 | Shinzato<br>Matz<br>Selmoni | 10,886,031<br>10,889,427<br>10,884,619 | 1 | LOC114969402 | ns | 10,884,051 | 10,891,914 | - |
| 61 | chr13<br>NC_058078.1 | 11,510,001-<br>11,510,001 | 0.063 | Selmoni<br>Torquato<br>Shinzato<br>Matz | 11,516,500<br>11,510,276<br>11,512,003<br>11,519,467 | 2 | LOC114966268<br>LOC114966267 | -<br>- | 11,511,156<br>11,505,833 | 11,517,786<br>11,511,217 | -<br>- |
| 62 | chr13<br>NC_058078.1 | 17,730,001-<br>17,730,001 | 0.068 | Selmoni<br>Torquato<br>Shinzato<br>Cooke | 17,734,360<br>17,736,884<br>17,731,410<br>17,733,000 | 1 | LOC114959494 | ? | 17,731,815 | 17,736,210 | - |
| 63 | chr14<br>NC_058079.1 | 8,230,001-<br>8,230,001 | 0.068 | Shinzato<br>Selmoni<br>Matz | 8,238,308<br>8,239,161<br>8,238,167 | 2 | LOC114962569<br>LOC114962580 | +<br>- | 8,237,242<br>8,223,640 | 8,290,054<br>8,231,522 | Rho GTPase-activating protein 29<br>Plasminogen activator inhibitor 1 R--binding protein |
| 64 | chr14<br>NC_058079.1 | 12,470,001-<br>12,470,001 | 0.068 | Drury<br>Cooke<br>Shinzato<br>Matz | 12,475,609<br>12,476,827<br>12,474,460<br>12,472,753 | NA | NA | ? | NA | NA | NA |
| 65 | chr14<br>NC_058079.1 | 16,280,001-<br>16,280,001 | 0.093 | Matz<br>Selmoni<br>Shinzato | 16,289,268<br>16,286,493<br>16,288,451 | 2 | LOC114956529<br>LOC114956528 | -<br>? | 16,285,987<br>16,271,252 | 16,307,073<br>16,285,140 | Afadin<br>- |
| 66 | NW_025322615.1 | 630,001-<br>630,001 | 0.096 | Shinzato<br>Matz<br>Selmoni | 636,150<br>635,134<br>636,149 | 1 | LOC114947105 | ns | 632,316 | 657,365 | Ephrin type-A receptor 3 |
| 67 | NW_025322620.1 | 180,001-<br>180,001 | 0.08 | Cooke<br>Torquato<br>Drury<br>Shinzato | 184,043<br>181,579<br>182,675<br>181,902 | 1 | LOC114967789 | ? | 180,442 | 182,029 | ATP-dependent D- helicase Q1 |
| 68 | NW_025322625.1 | 1,180,001-<br>1,180,001 | 0.061 | Cooke<br>Shinzato<br>Torquato | 1,183,313<br>1,182,723<br>1,186,306 | 1 | LOC114963338 | + | 1,179,631 | 1,219,065 | Oncoprotein-induced transcript 3 protein |
| 69 | NW_025322625.1 | 1,210,001-<br>1,210,001 | 0.068 | Torquato<br>Drury<br>Shinzato<br>Cooke | 1,218,994<br>1,216,031<br>1,219,814<br>1,211,730 | 1 | LOC114963338 | + | 1,179,631 | 1,219,065 | Oncoprotein-induced transcript 3 protein |

| | Genomic windows | | | SNPs | | Genes | | | | | |
|---|---|---|---|---|---|---|---|---|---|---|---|
| | Chromosome/ contig | Start- End | q | Datasets | Position | # | ID | DGE | Start | End | Annotated Protein |
| 70 | NW_025322626.1 | 530,001-<br>530,001 | 0.032 | Cooke<br>Torquato<br>Matz<br>Shinzato | 539,776<br>536,316<br>539,570<br>535,429 | 3 | LOC114967317<br>LOC114973321<br>LOC114967303 | ns<br>ns<br>ns | 530,249<br>539,254<br>519,578 | 532,042<br>543,469<br>530,043 | -<br>TNF receptor-associated factor 6<br>- |
| 71 | NW_025322627.1 | 930,001-<br>930,001 | 0.091 | Torquato<br>Shinzato<br>Cooke<br>Selmoni | 932,032<br>932,783<br>931,643<br>931,999 | 1 | LOC122949945 | + | 930,352 | 941,249 | Receptor-type tyrosine-protein phosphatase delta |
| 72 | NW_025322628.1 | 660,001-<br>660,001 | 0.055 | Torquato<br>Shinzato<br>Matz | 663,377<br>667,339<br>668,559 | 2 | Trnat-ggu-48<br>LOC114964724 | ?<br>- | 661,667<br>669,364 | 661,739<br>671,368 | -<br>Collagen alpha-3(VI) chain |
| 73 | NW_025322633.1 | 40,001-<br>40,001 | 0.096 | Matz<br>Shinzato<br>Cooke<br>Drury | 43,357<br>43,678<br>41,589<br>43,568 | 1 | LOC122950252 | + | 38,861 | 41,410 | - |
| 74 | NW_025322636.1 | 140,001-<br>140,001 | 0.086 | Torquato<br>Cooke<br>Selmoni<br>Matz<br>Shinzato | 141,624<br>147,245<br>141,534<br>146,878<br>141,524 | 1 | LOC114956952 | ns | 135,597 | 162,606 | NFX1-type zinc finger-containing protein 1 |
| 75 | NW_025322654.1 | 120,001-<br>120,001 | 0.055 | Matz<br>Shinzato<br>Cooke | 121,009<br>120,478<br>121,157 | 1 | LOC122951140 | ? | 128,457 | 130,976 | Zinc finger protein 862 |
| 76 | NW_025322656.1 | 120,001-<br>120,001 | 0.082 | Torquato<br>Selmoni<br>Cooke | 120,311<br>120,414<br>126,380 | 2 | LOC114969246<br>LOC114973944 | ?<br>ns | 117,351<br>73,292 | 121,106<br>156,869 | -<br>MAM and LDL-receptor class A domain-containing protein 1 |
| 77 | NW_025322657.1 | 200,001-<br>200,001 | 0.055 | Shinzato<br>Torquato<br>Drury | 201,932<br>203,463<br>203,486 | 1 | LOC122951222 | ? | 203,060 | 203,838 | Retrovirus-related Pol polyprotein from transposon 412 |
| 78 | NW_025322668.1 | 270,001-<br>270,001 | 0.099 | Cooke<br>Selmoni<br>Matz<br>Torquato | 276,644<br>277,190<br>276,239<br>277,146 | NA | NA | ? | NA | NA | NA |
| 79 | NW_025322695.1 | 70,001-<br>70,001 | 0.091 | Matz<br>Cooke<br>Torquato | 72,978<br>77,141<br>74,976 | 1 | LOC114954361 | ? | 70,393 | 72,360 | Melanocyte-stimulating hormone receptor |
| 80 | NW_025322750.1 | 50,001-<br>50,001 | 0.068 | Cooke<br>Matz<br>Torquato | 52,775<br>54,970<br>54,239 | 1 | LOC114972120 | ? | 56,285 | 57,589 | - |
| 81 | NW_025322770.1 | 150,001-<br>150,001 | 0.068 | Selmoni<br>Cooke<br>Matz | 152,302<br>150,959<br>155,219 | 2 | LOC114965893<br>LOC114965872 | ns<br>? | 154,491<br>149,693 | 163,294<br>153,341 | -<br>- |
| 82 | NW_025322777.1 | 390,001-<br>390,001 | 0.041 | Cooke<br>Matz<br>Torquato | 397,719<br>397,353<br>397,693 | 2 | LOC122953434<br>LOC114947948 | ?<br>+ | 399,154<br>377,872 | 400,002<br>396,917 | -<br>Hemicentin-1 |
| 83 | NW_025322890.1 | 20,001-<br>20,001 | 0.055 | Matz<br>Torquato<br>Cooke | 25,314<br>25,035<br>24,868 | 2 | LOC114967591<br>LOC122954621 | ?<br>? | 24,130<br>27,134 | 26,945<br>29,277 | -<br>- |
| 84 | NW_025323125.1 | 230,001-<br>230,001 | 0.032 | Selmoni<br>Cooke<br>Torquato<br>Matz | 237,886<br>237,592<br>237,862<br>237,375 | NA | NA | ? | NA | NA | NA |
| 85 | NW_025323127.1 | 80,001-<br>80,001 | 0.069 | Cooke<br>Torquato<br>Selmoni | 83,352<br>85,647<br>85,600 | NA | NA | ? | NA | NA | NA |

**Table S4. Gene ontology enrichment analysis.**

Shown are the eleven gene ontology (GO) terms found as enriched in the 85 genomic regions where adaptive signals overlapped from different datasets. For every GO term, the table shows the ID, the description, the number of genes with the annotation across the genome, the setRank statistics of enrichment, and the p-value and adjusted p-value of the enrichment test.

| Gene Ontology ID | Term description | Size | setRank | P-value | Adj P-value |
|---|---|---|---|---|---|
| GO:0016779 | nucleotidyltransferase activity | 30 | 0.04718 | 2.50E-05 | 0.000274 |
| GO:0031072 | heat shock protein binding | 38 | 0.087284 | 1.31E-04 | 0.001315 |
| GO:0051082 | unfolded protein binding | 93 | 0.04718 | 1.31E-04 | 0.001315 |
| GO:0001640 | adenylate cyclase inhibiting G protein-coupled glutamate receptor activity | 15 | 0.15962 | 5.06E-04 | 0.003539 |
| GO:0099507 | ligand-gated ion channel activity involved in regulation of presynaptic membrane potential | 10 | 0.04718 | 3.93E-04 | 0.003539 |
| GO:0015277 | kainate selective glutamate receptor activity | 12 | 0.060548 | 4.49E-04 | 0.003539 |
| GO:0008066 | glutamate receptor activity | 16 | 0.086281 | 5.06E-04 | 0.003539 |
| GO:0008934 | inositol monophosphate 1-phosphatase activity | 3 | 0.087284 | 7.30E-04 | 0.005842 |
| GO:0043812 | phosphatidylinositol-4-phosphate phosphatase activity | 4 | 0.04718 | 7.30E-04 | 0.005842 |
| GO:0004051 | arachidonate 5-lipoxygenase activity | 3 | 0.04718 | 1.35E-03 | 0.009434 |

**Table S5. Coefficients of *Acropora* cover change models.**

Shown are the coefficients of models describing changes in *Acropora* cover in the North Great Barrier Reef during the 2016 and 2017 heat waves (*38*). For each model shown is the response variable, the fixed effects—with corresponding estimate, standard error, z-value and p-value—and the random factors. (A) shows the coefficients for a model accounting for heat intensity during the heatwave, (B) for a model accounting for heat intensity and the expected frequency of heat-adaptive genotypes, (C) for a model accounting for heat intensity and past heat exposure.

| **A - Max DHW between surveys** | | | | | | | |
|---|---|---|---|---|---|---|---|
| **Response variable** | **Fixed effects** | | | | | **Random factors** | |
| | ***Coefficient*** | ***Estimate*** | ***Std. Error*** | ***z value*** | ***P-value*** | ***Factor*** | ***St. Dev*** |
| Relative *Acropora* cover change between two surveys on different years | (Intercept) | -0.02 | 0.01 | -1.44 | 0.1489 | Reef of survey (intercept) | 0.011 |
| | Max DHW between surveys | 0 | 0 | -1.18 | 0.2389 | Marine province (intercept) | 0.023 |
| | | | | | | Residuals | 0.036 |

| **B - Max DHW between surveys x historical maximal DHW** | | | | | | | |
|---|---|---|---|---|---|---|---|
| **Response variable** | **Fixed effects** | | | | | **Random factors** | |
| | ***Coefficient*** | ***Estimate*** | ***Std. Error*** | ***z value*** | ***P-value*** | ***Factor*** | ***St. Dev*** |
| Relative *Acropora* cover change between two surveys on different years | (Intercept) | 0.01 | 0.04 | 0.21 | 0.834 | | |
| | Max DHW between surveys | -0.01 | 0 | -2.35 | 0.0187 | Reef of survey (intercept) | 5e-06 |
| | Exp. freq. Adaptive GT | -0.01 | 0.01 | -0.79 | 0.4288 | Marine province (intercept) | 0.021 |
| | Interaction: Max DHW*Exp. Fr. Ad. GT | 0 | 0 | 2.05 | 0.0407 | Residuals | 0.036 |

| **C - Max DHW between surveys x Expected freq. Heat-adaptive genotypes** | | | | | | | |
|---|---|---|---|---|---|---|---|
| **Response variable** | **Fixed effects** | | | | | **Random factors** | |
| | ***Coefficient*** | ***Estimate*** | ***Std. Error*** | ***z value*** | ***P-value*** | ***Factor*** | ***St. Dev*** |
| Relative *Acropora* cover change between two surveys on different years | (Intercept) | 0.12 | 0.03 | 4.05 | 5.17e-05 | | |
| | Max DHW between surveys | -0.027 | 0.01 | -4.27 | 1.95e-05 | Reef of survey (intercept) | 0.009 |
| | Exp. freq. Adaptive GT | -0.25 | 0.05 | -4.45 | 8.70e-06 | Marine province (intercept) | 0.007 |
| | Interaction: Max DHW*Exp. Fr. Ad. GT | 0.042 | 0.01 | 4.03 | 5.65e-05 | Residuals | 0.036 |

**Table S6. Parameters of Mutations-Area Relationship Linear Mixed Model.**

Shown are the parameters of the linear mixed models explaining variation in log-transformed mutations (logM) across spatial subsamples from the *Acropora* datasets. In every model (rows), different fixed (logA: log-transformed area of the subsample, logSD: log-transformed mean sequencing depth of samples in the subsample) and random effects were used. The deviance information criterion (DIC) indicates the model's goodness of fit (lower values mean higher fit).

| MID | Response variable | Fixed variables | Random effects | DIC |
|---|---|---|---|---|
| 1 | logM | - | Dataset | 440 |
| 2 | logM | - | Dataset + Species | 440 |
| 3 | logM | - | Dataset + GenomicSubSample | 377 |
| 4 | logM | - | Dataset + GenomicSubSample + Spatial Index | -2047 |
| 5 | logM | logA | Dataset + GenomicSubSample + Spatial Index | -3508 |
| 6 | logM | logA + logSD | Dataset + GenomicSubSample + Spatial Index | -4282 |

### Supplementary Figures

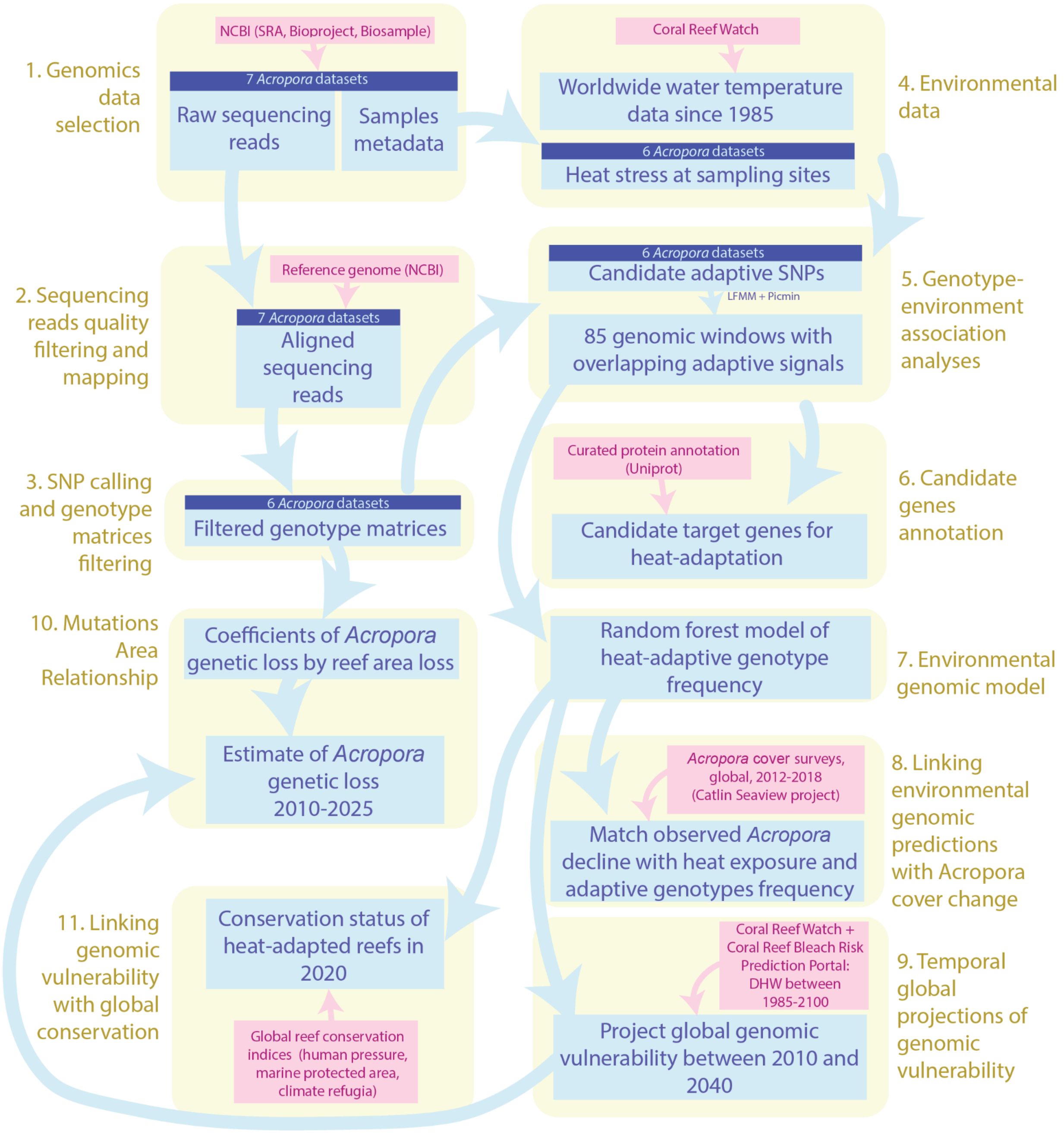


**Figure S1. Workflow of the study.**

Shown in yellow are the different steps of the study, in red the input from external data sources for the analysis, and in blue the analytical inputs and outputs from the different steps.

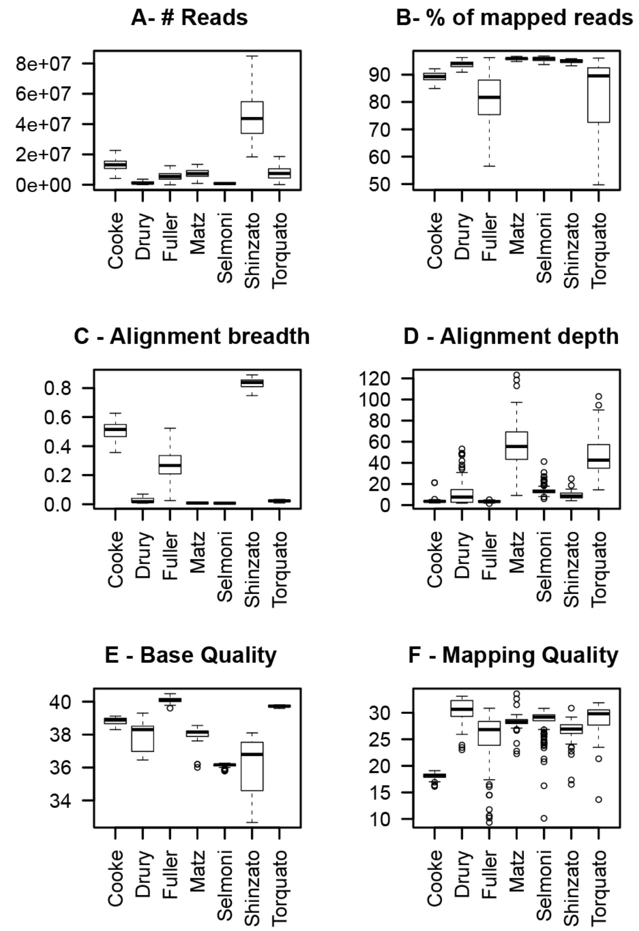


**Figure S2. Sequencing read mapping stats.**

Shown are the sequencing and alignment statistics by dataset: (A) distribution of number of reads across samples, (B) percentage of reads successfully mapped to the reference across samples, (C) genome breadth covered by alignment across samples, (D) mean sequencing depth across samples, (E) mean base quality scores across samples, (F) mean quality of read-to-reference mapping across samples.

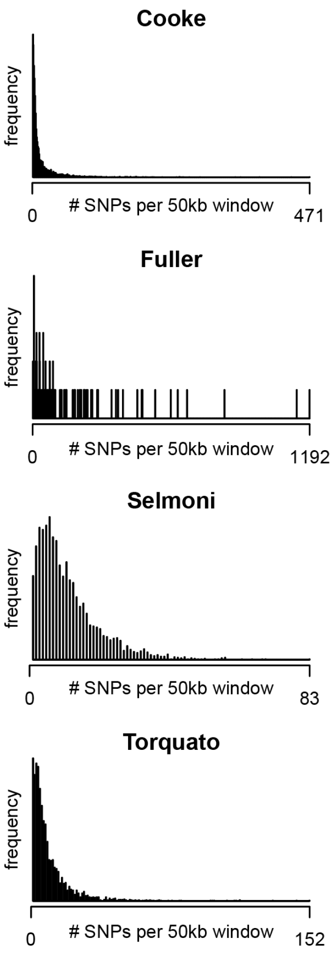


**Figure S3. Distribution of SNPs across genomes.**

Shown are the number of single nucleotide polymorphisms (SNPs) called across 50 kbs genomic windows in the seven *Acropora* datasets.

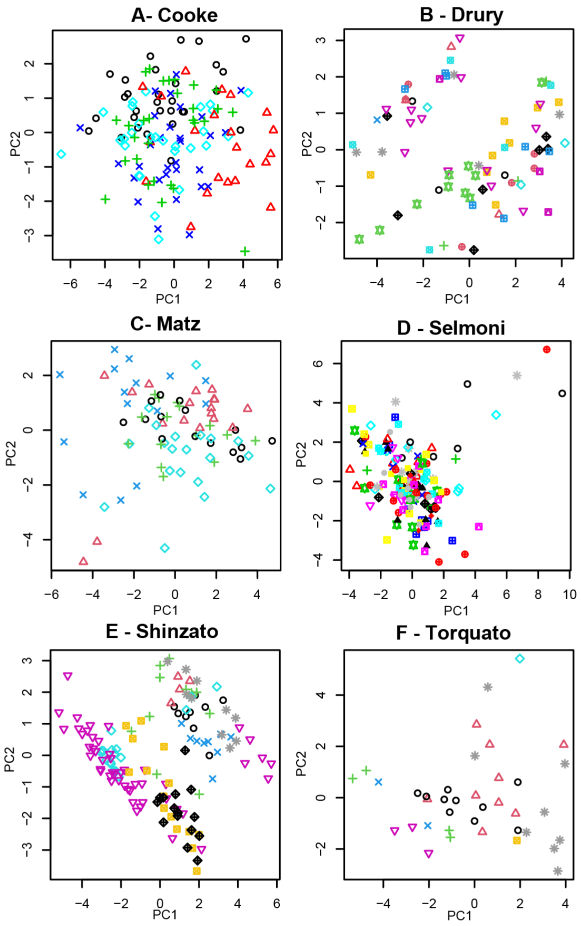


**Figure S4. Principal component analysis of sequencing and alignment statistics.**

Shown are the main two axes of variation of sequencing and alignment statistics across samples from the six *Acropora* datasets. The axes of variation were computed by running a Principal Component Analysis on by-sample sequencing and alignment statistics: number of reads, percentage of mapped reads, alignment breadth, alignment depth, mean base quality, mean mapping quality. Colored symbols correspond to reefs where samples were collected from.

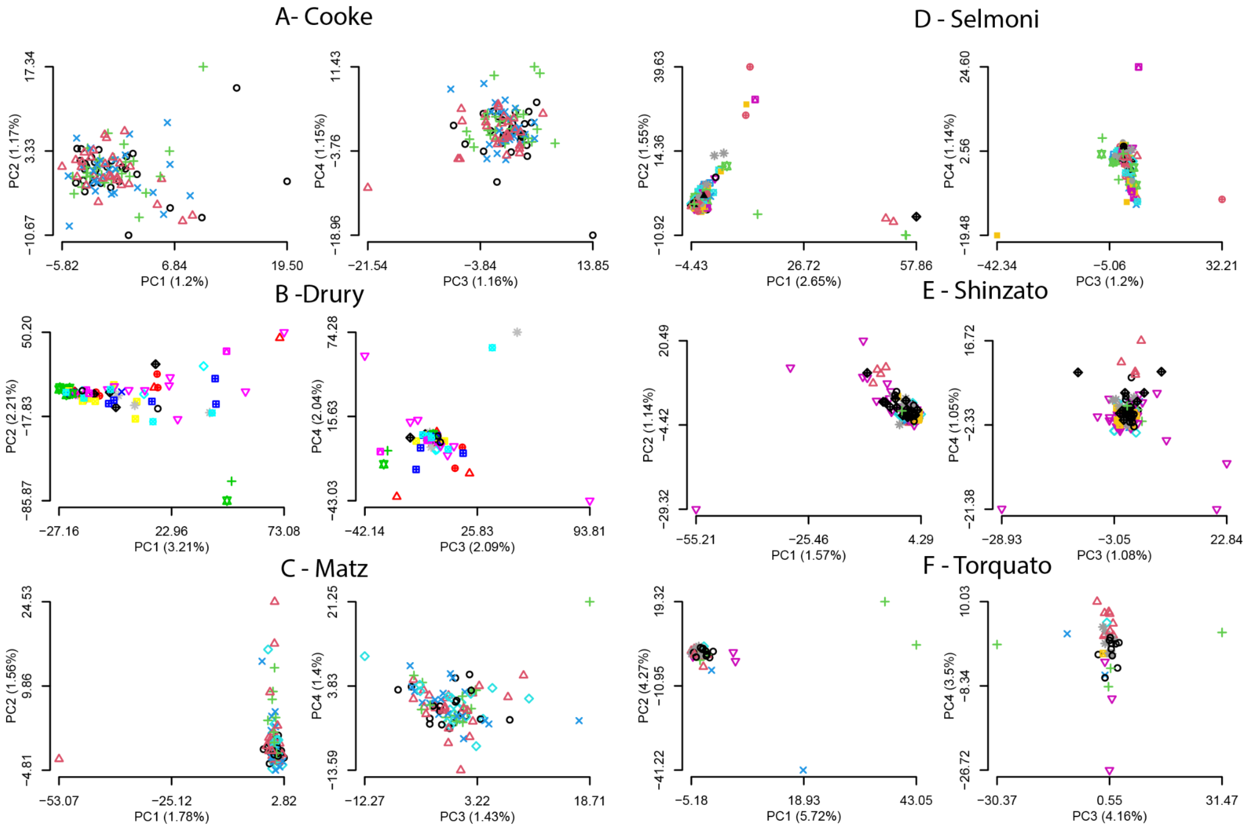


**Figure S5. Principal component analysis of genotype matrices.**

Shown are the main four axes of genetic variation across samples from the six *Acropora* datasets. The axes of variation were computed by running a Principal Component Analysis on the genotype matrix of every dataset. Colored symbols correspond to distinct sampling locations.

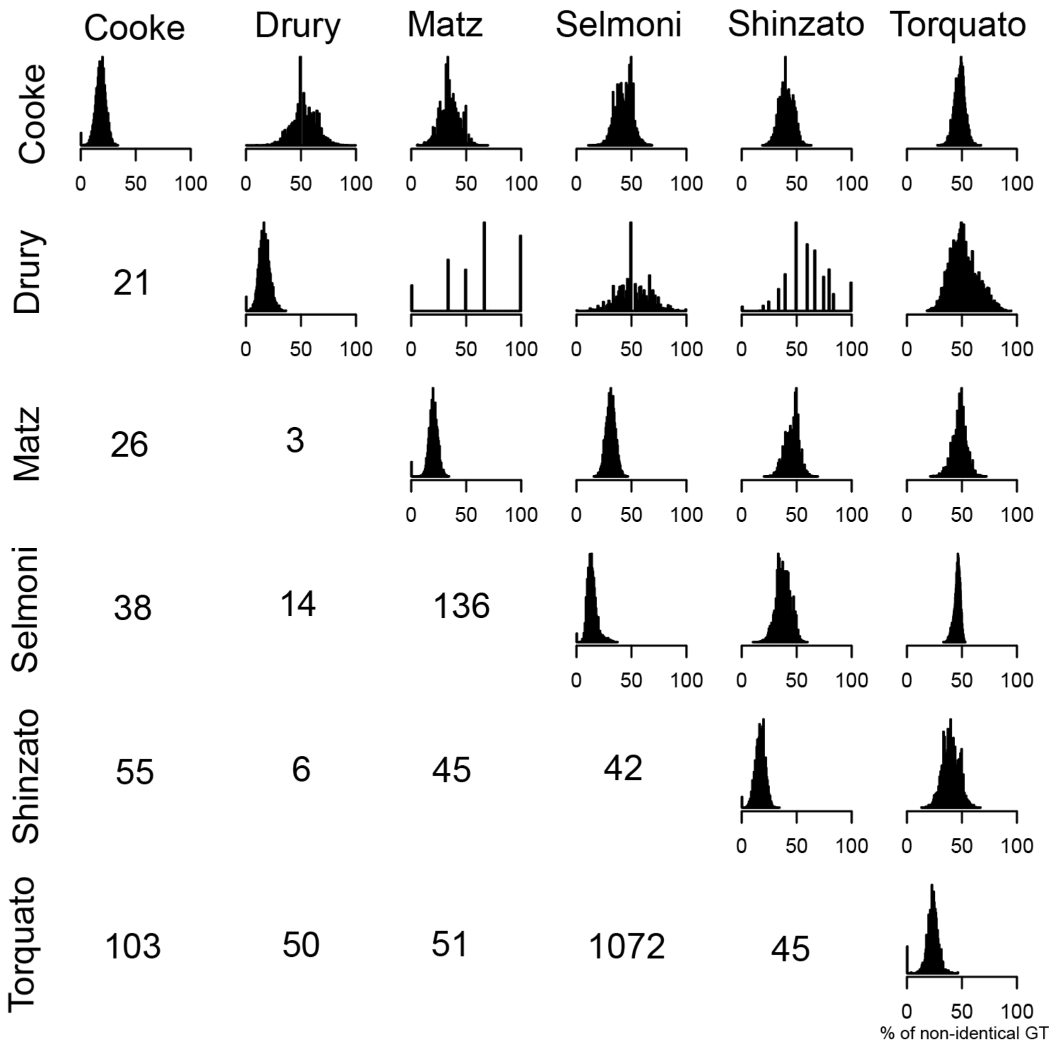


**Figure S6. Shared SNPs and % of non-identical genotypes across datasets.**

Below the diagonal are shown the number of single nucleotide polymorphisms shared between pairs of datasets. Focusing on these SNPs, the percentage of non-identical genotypes - *i.e.* the genetic distance - was calculated between pairs of samples for different datasets. Above the diagonal are shown the distribution of genetic distance between datasets, the diagonal shows the distribution of genetic distance within datasets.

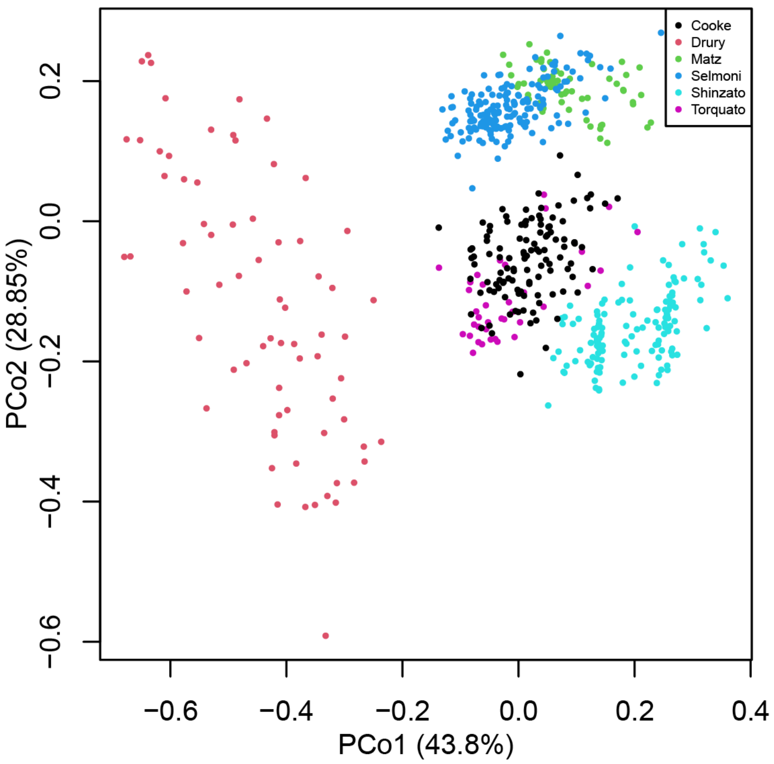


**Figure S7. Genetic proximity of all samples from the *Acropora* datasets.**

Shown are the main two axes summarizing the genetic distances between the six *Acropora* datasets (colors). The axes were computed by running a Principal Coordinate Analysis on the matrix of genetic distance between all samples, computed as the percentage of non-identical genotypes across the shared single nucleotide polymorphisms.

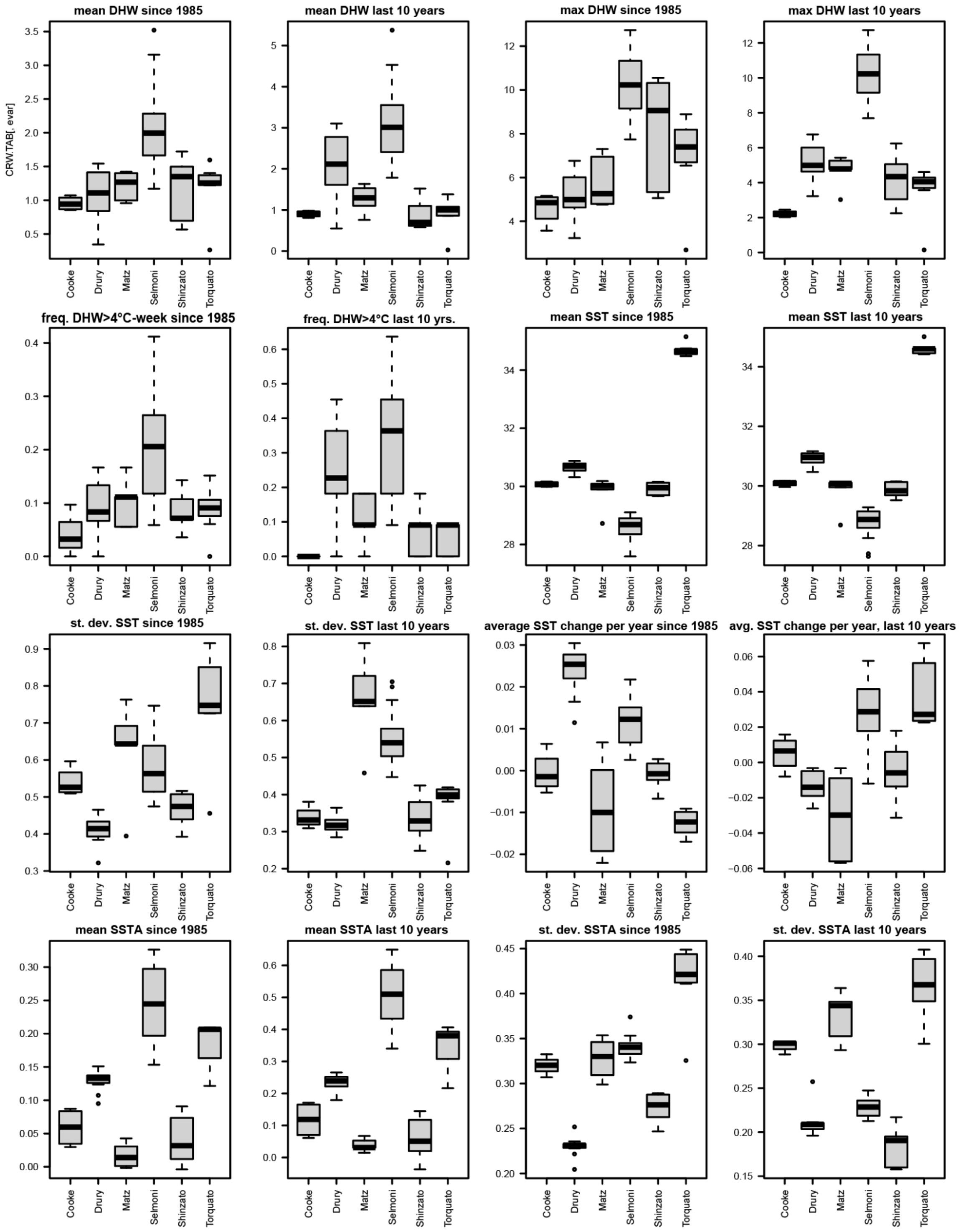


**Figure S8. Heat stress variables distribution across datasets.**

For the sampling sites of the *Acropora* genomic datasets, shown are the distribution of 16 heat stress variables describing temporal trends of annual maximal Sea Surface Temperature (SST), annual maximal Degree Heating Week (DHW), and annual mean Sea Surface Temperature Anomaly (SSTA). Eight variables focus on temporal trends from 1985 until the year of sampling, and eight variables for the ten years before sampling.

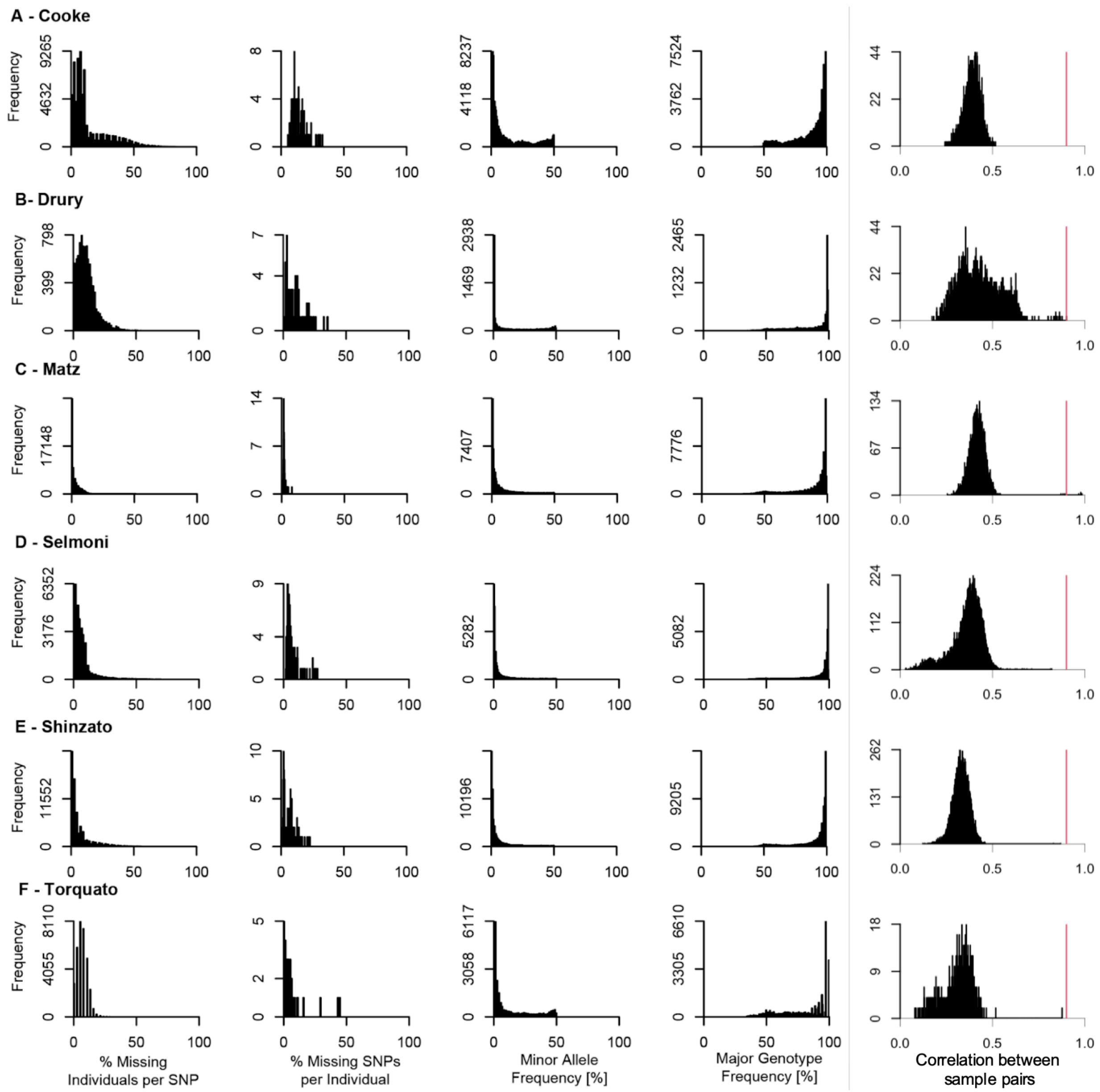


**Figure S9. Missing values, allele frequencies statistics and allele frequency correlation**

Shown are, for every *Acropora* genomics dataset (A-F), the following statistics of the genotype matrix: percentage of missing individuals per single nucleotide polymorphism (SNP), percentage of missing SNP per individual, minor allele frequency per SNP, major genotype frequency per SNP, and the allele frequency correlation across samples (vertical red line represents the cutoff thresholds for clones R=0.9).

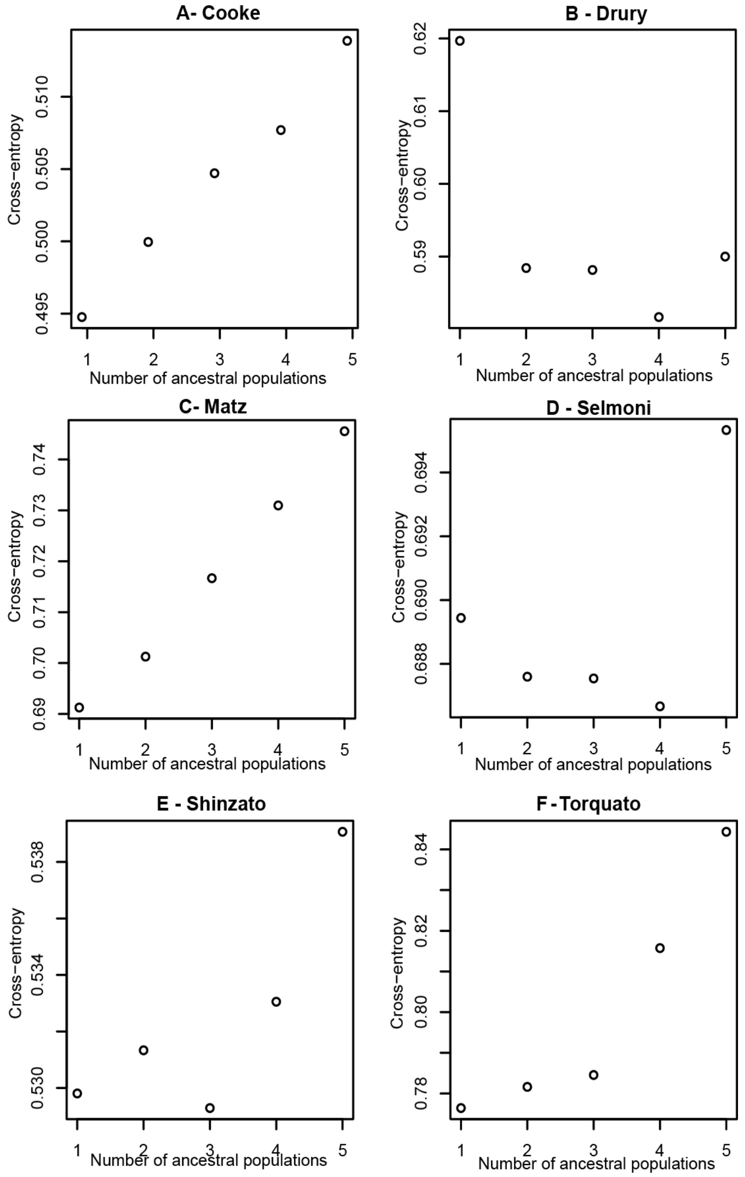


**Figure S10. Cross-entropy estimation of the number of ancestral populations.**

Shown are the cross-entropy values for different numbers of ancestral populations across the different *Acropora* datasets (A-F). Lower cross-entropy indicates higher goodness of fit of the number of ancestral populations to the genetic diversity observed in the genotype matrix.

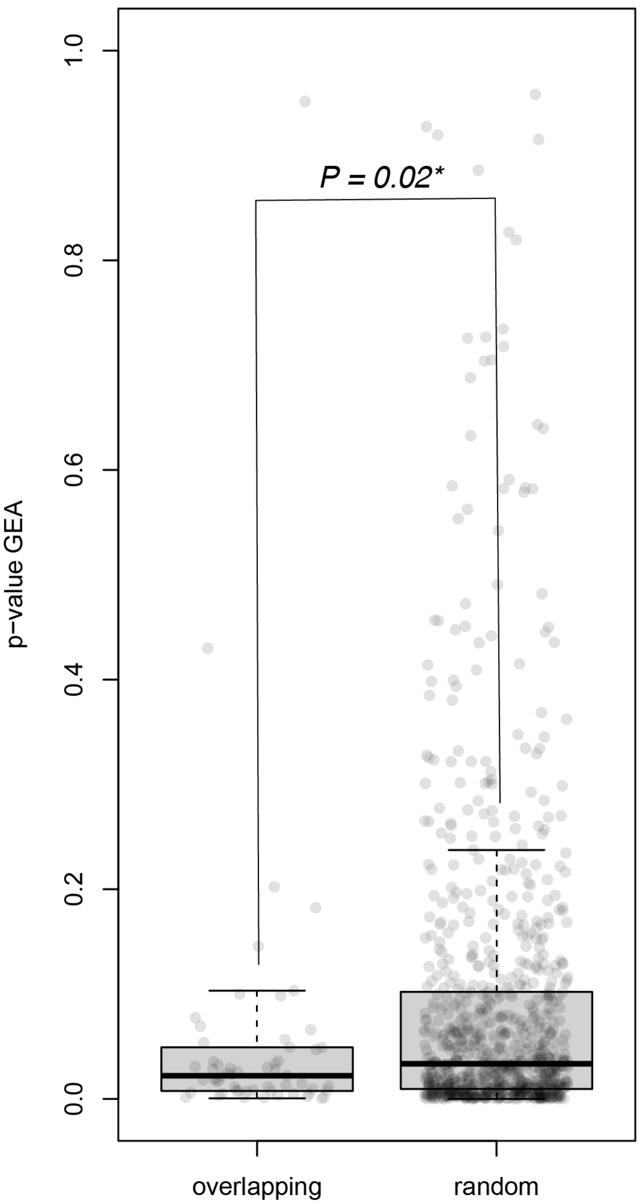


**Figure S11. Validation of overlapping adaptive signals on an independent *Acropora* dataset.**

Shown are the genome-wide p-values describing the genotype-environment association (GEA) between DHW-max and SNPs from an *A. millepora* population sampled on the central GBR in 2017 (*1*). Every point corresponds to the lowest GEA p-value found within a 10 kbs genomic window. On the left side: p-value distributions for 85 genomic windows showing significant overlap of heat-adaptive signals in three or more *Acropora* populations from the overlap analysis. On the right side: p-value distribution for 1000 random genomic windows. The distributions of p-values between overlapping and random genomic window were compared using a Wilcoxon rank sum test (test P-value shown on top).

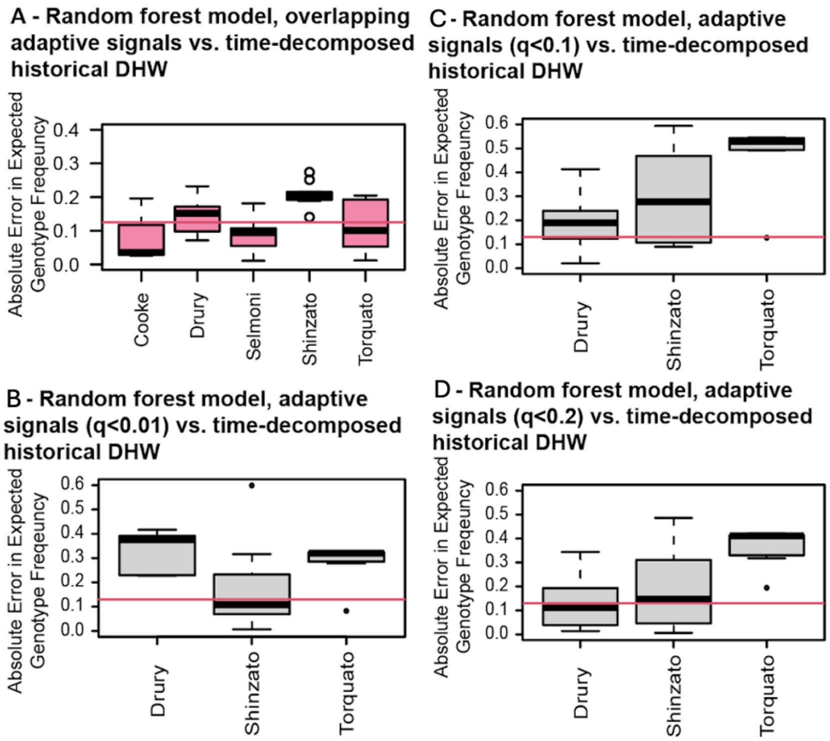


**Figure S12. Cross validation of the environmental genomic model.**

Shown are the results of the cross validation assessing the predictive power of different environmental genomic models. All models predict the frequency of candidate adaptive genotypes at a given reef from historical Degree Heating Week trends (DHW). For each model, a leave-one-dataset-out cross validation approach was used to calculate absolute errors in predicting the expected genotype frequencies.
In all models, historical maximal DHW was decomposed in 5-year windows (0 to 5, 5 to 10, 10 to 15, 15 to 20 and 20 to 25 years before sampling), and a random forest model estimated genotype frequencies from decomposed DHW trends. In (A), the model is based on candidate adaptive genotypes detected in the same genomic window in at least three datasets (FDR<0.1). In the other panels, models are based on candidate adaptive genotypes detected in any dataset (without overlap analysis) using different FDR thresholds (B = 0.01, C = 0.1, D = 0.2). The red horizontal line is a reference to the mean absolute error in (A).

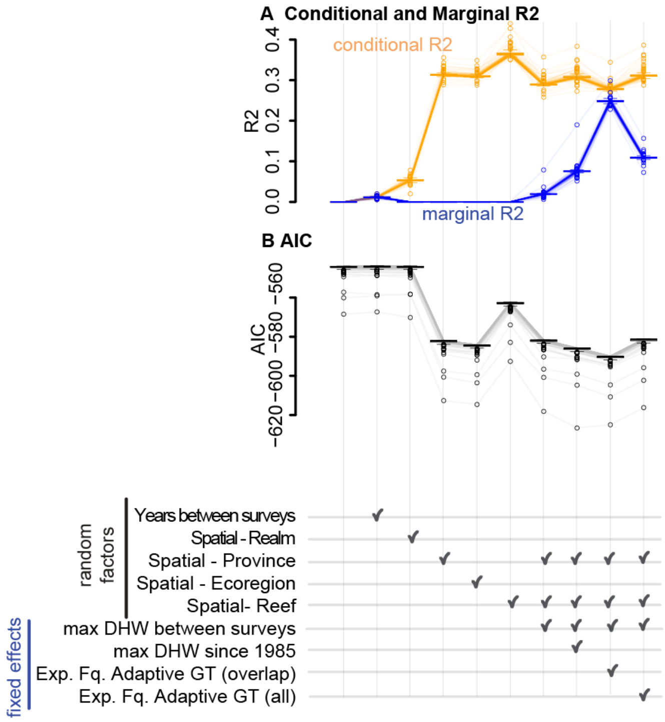


**Figure S13. Performance of models explaining *Acropora* cover decline across Indo-Pacific.**

(A) shows the distribution of the conditional (orange) and marginal (blue) coefficient of determination (R2) for models using different explanatory variables to describe *Acropora* cover change across the Indo-Pacific. Conditional R2 represents the % of variance explained by the entire model, marginal R2 the % explained by fixed effects. The models use different combinations of explanatory variables: number of years between coral cover surveys, spatial autocorrelation between survey sites at realm, province, ecoregion and reef level, maximal Degree Heating Week (DHW) measured between the dates of the field surveys, the maximal DHW since 1985, and the expected frequency of adaptive genotypes based on an environmental genomic model. The environmental genomic model can be based on candidate adaptive genotypes found in the same genomic regions in different datasets (overlap), or any candidate adaptive genotype found in any dataset (all). The lines on the background represent the conditional R2 estimates obtained using a jackknife resampling approach. (B) shows the distribution of the Akaike Information Criterion (AIC) for the same models.

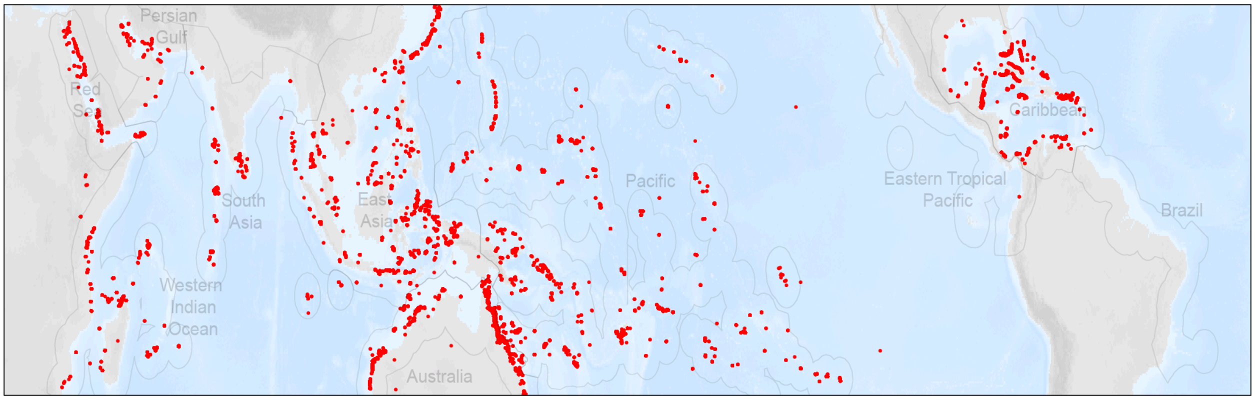


**Figure S14. Worldwide distribution of Acorpora records from the Ocean Biodiversity Information System (OBIS).**

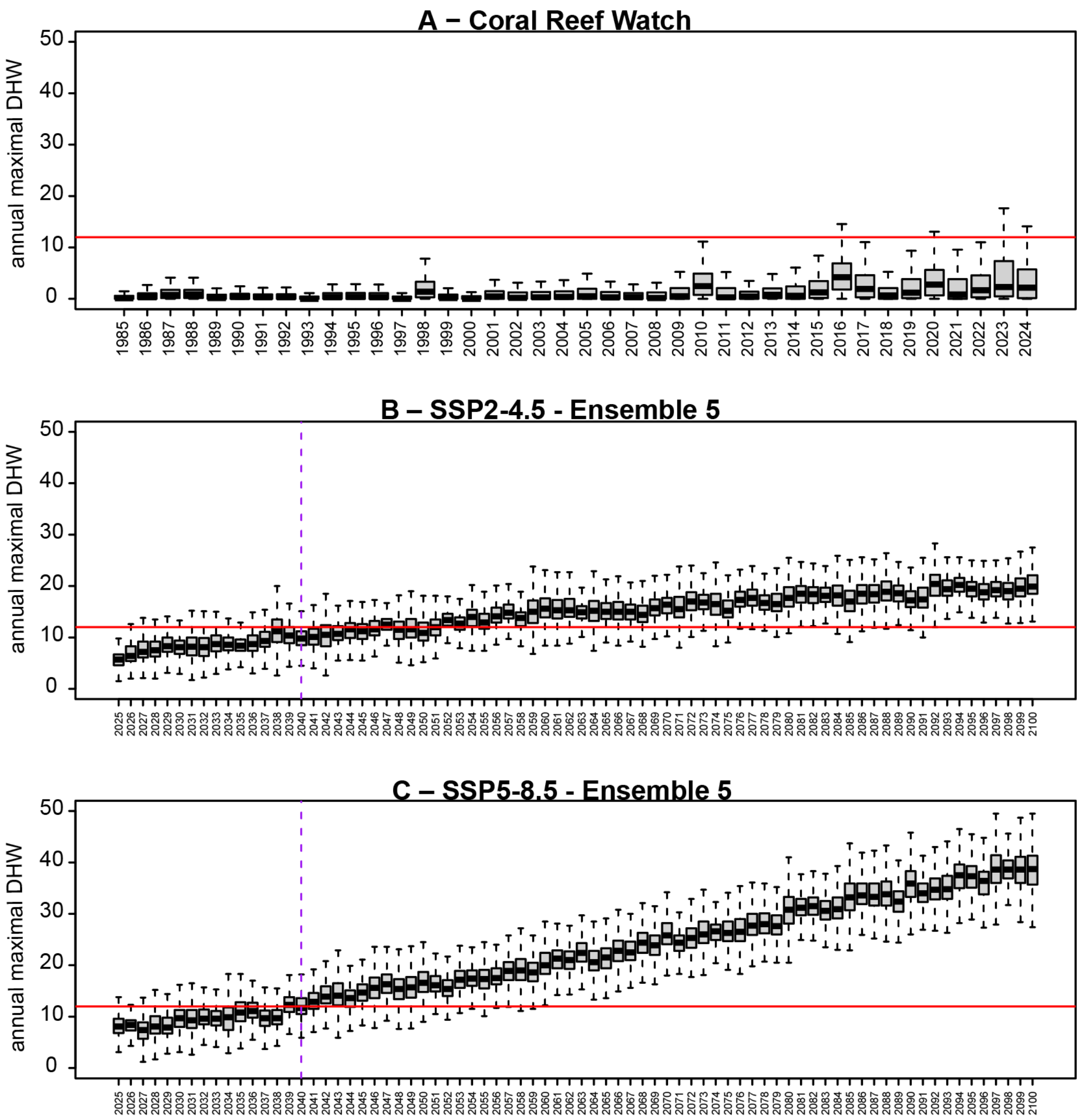


**Figure S15. Measured and projected maximal annual Degree Heating Week.**

Shown are the year-by-year distribution of measured (A) and projected (B, C) values of maximal Degree Heating Week (DHW) across reefs worldwide. Distributions in (A) are based contemporary measurements (1985-2024) from the Coral Reef Watch, whereas distributions in (B) and (C) from the Coral Reef Bleach Risk Prediction Portal (https://coralbleachrisk.net/), using the Ensemble 5 projections under SSP2-4.5 and SSP5-8.5 scenarios, respectively. The red horizontal line indicates the upper limit of the maximal DHW range used to build the environmental genomic model (~ 12°C-week). After 2040 (purple line), most of the reefs of the world are expected to be exposed to more than 12°C-week per year.

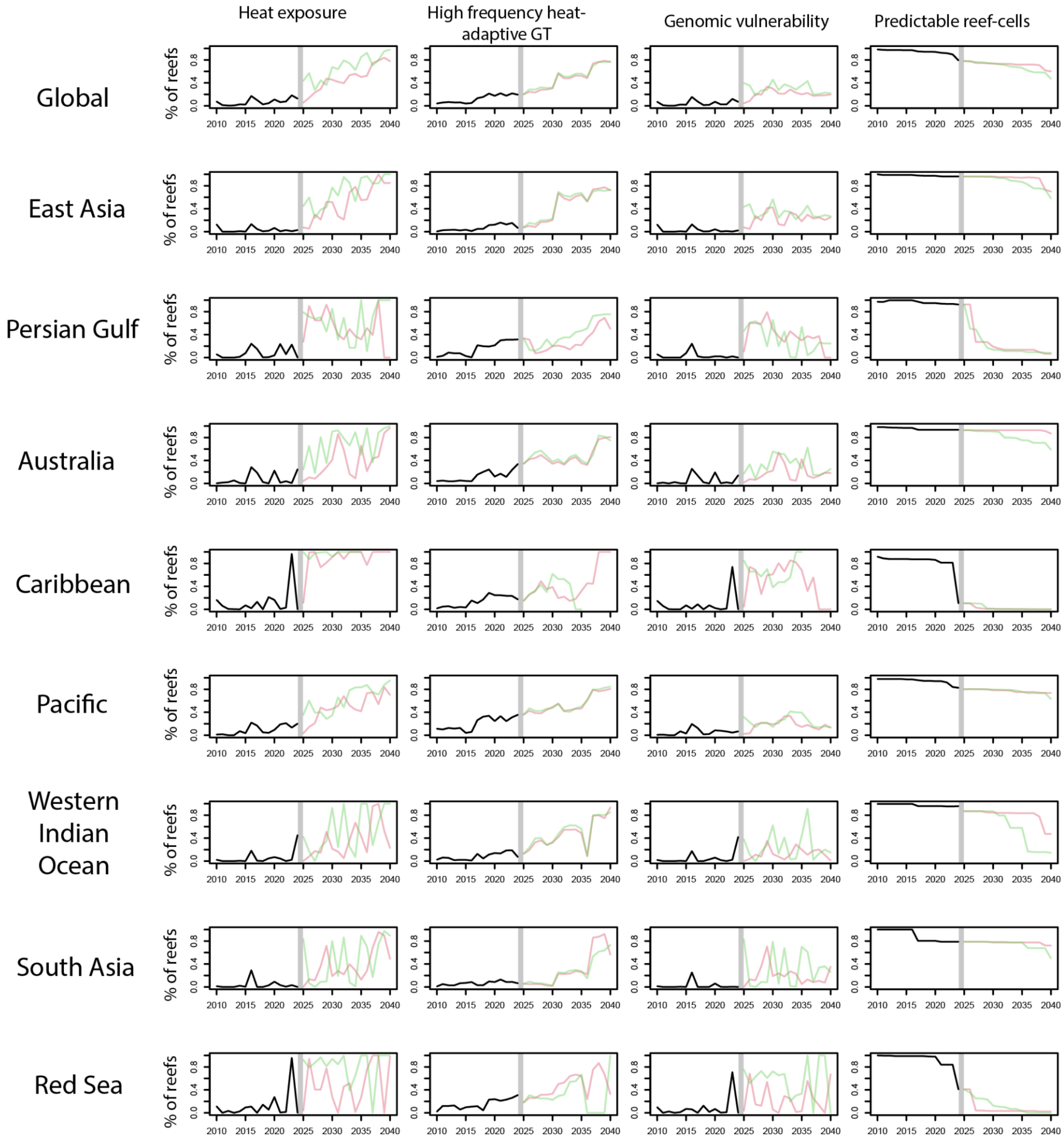


**Figure S16. Temporal projections of heat exposure, frequency of heat-adapted genotypes and genomic vulnerability.**

Shown are global (first row) and regional (the remaining rows) year-by-year projections between 2010 and 2040. The first column shows the year-by-year fraction of reefs exposed to a severe heatwave (*i.e.*, reefs exposed to a Degree Heating Week-DHW>8°C-week). The second column shows the year-by-year fraction of reefs with a high expected frequency of heat-adaptive genotypes (>70%), based on the environmental genomic model. The third column shows the year-by-year fraction of reefs under genomic vulnerability (*i.e.*, reefs with an expected frequency of heat-adaptive genotype <70%, exposed to DHW>8°C). The fourth column shows the year-by-year fraction of reefs that were usable for calculating the projections (*i.e.*, reefs with thermal history not exceeding the model predictive range). Projections before 2025 are based on the Coral Reef Watch data, projections after 2025 are based on the Coral Reef Bleach Risk Prediction Portal (https://coralbleachrisk.net/), using the Ensemble 5 projections under SSP2-4.5 (red line) and SSP5-8.5 (green line) scenarios.

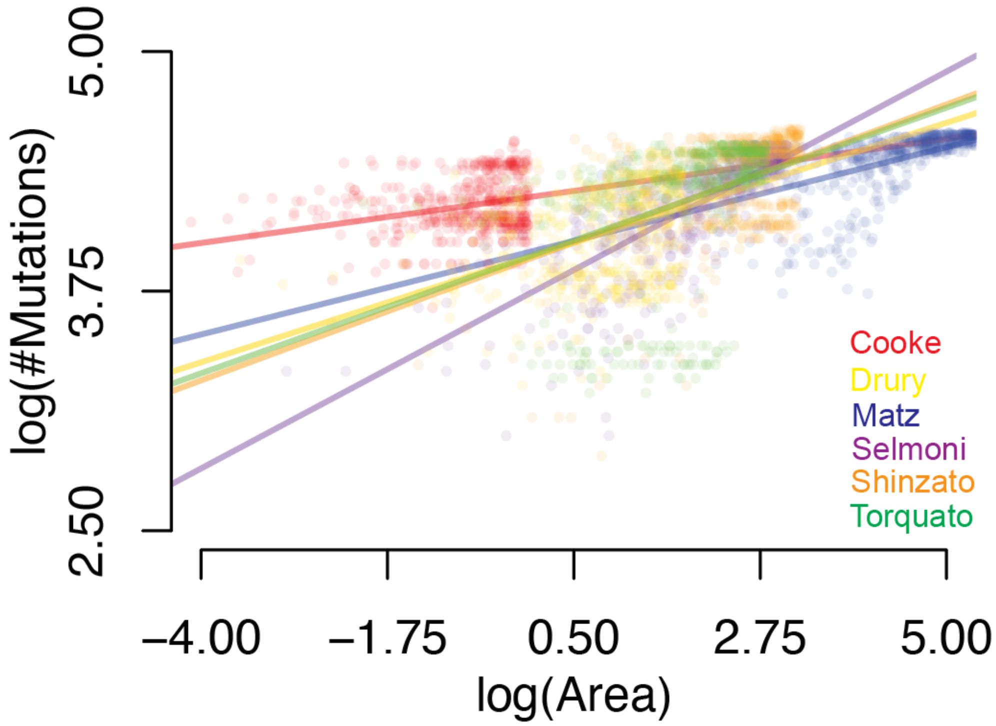


**Figure S17.** ***Acropora*** **mutations-area relationships (MAR).**

For each dataset (colors), shown are 500 spatial subsamples of the study region with (x-axis) the log-transformed area covered by the spatial subsample, and (y-axis) the log-transformed number of mutations across colonies in the subsampled area. Regression lines correspond to the z-mar parameter of every dataset.

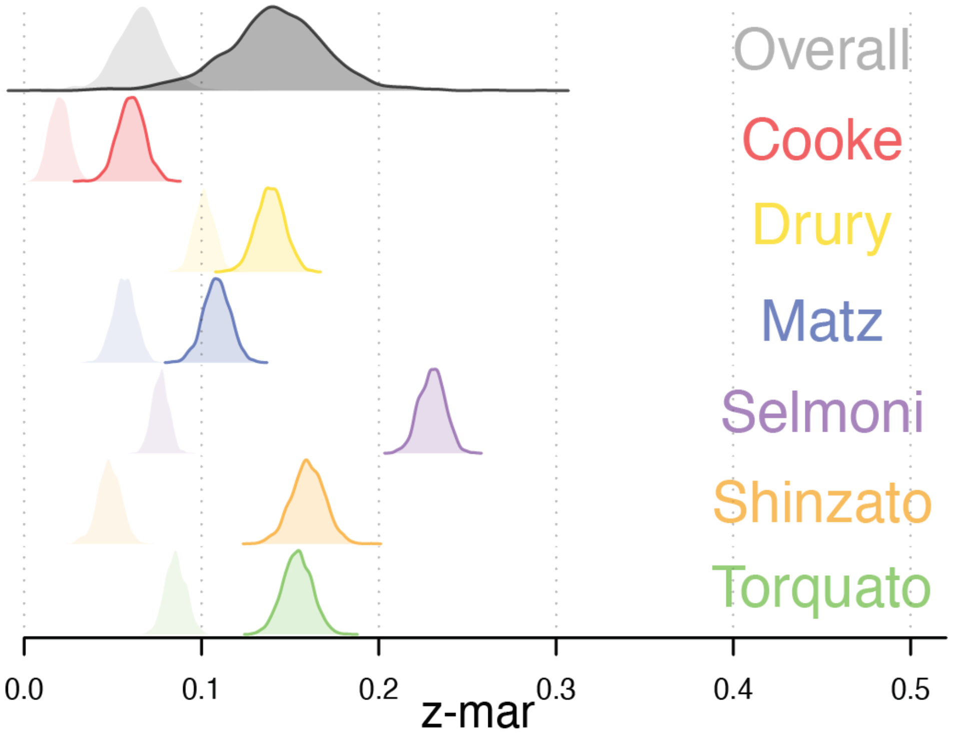


**Figure S18.** ***Acropora*** **z-mar coefficients.**

Shown are the z-mar coefficients for the six *Acropora* datasets (colored curves), and the overall average *Acropora* z-mar (black curve). The z-mar coefficients were calculated as the regression coefficients of the log-transformed area to the log-transformed number of mutations across datasets. The light-shaded curves represent the z-mar expectations under panmixia.

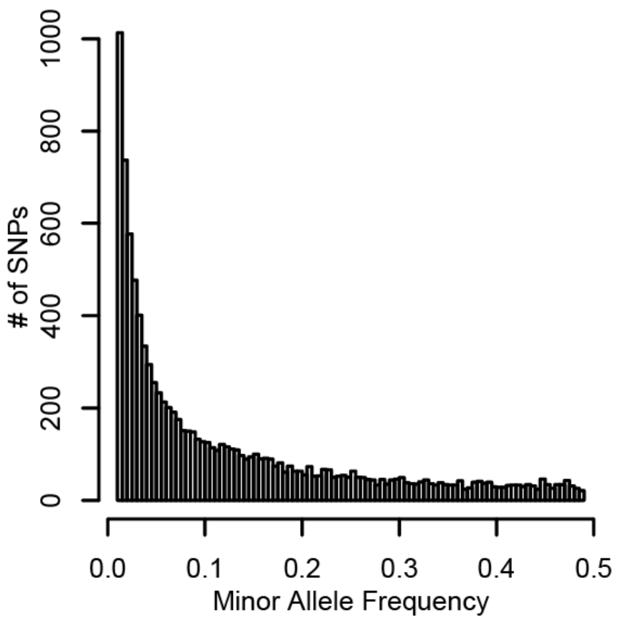


**Figure S19. Theoretical Allele Frequency Spectrum for a panmictic population.**

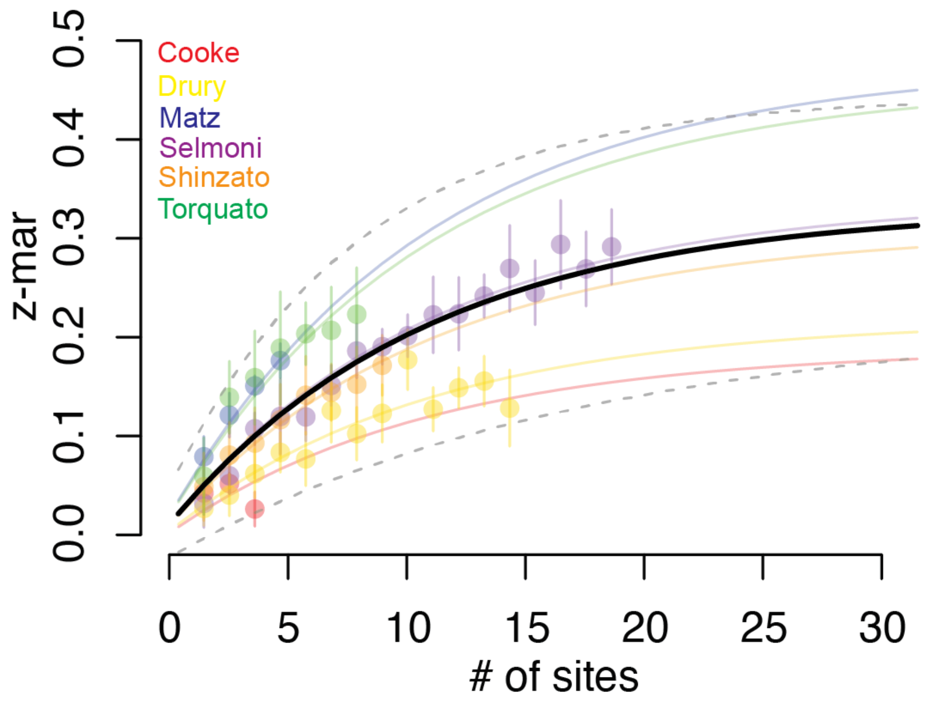


**Figure S20. Z-mar coefficient variation by limiting the number of sampling sites.**

For every *Acropora* dataset (colors), shown are the z-mar coefficients obtained while limiting the number of sampling sites in the Mutations Area Relationship calculation. The regression lines represent saturation curves of the number of sites-to-z-mar association for every dataset, and the black line represents the overall *Acropora* saturation curve (with intervals of confidence as dotted lines).

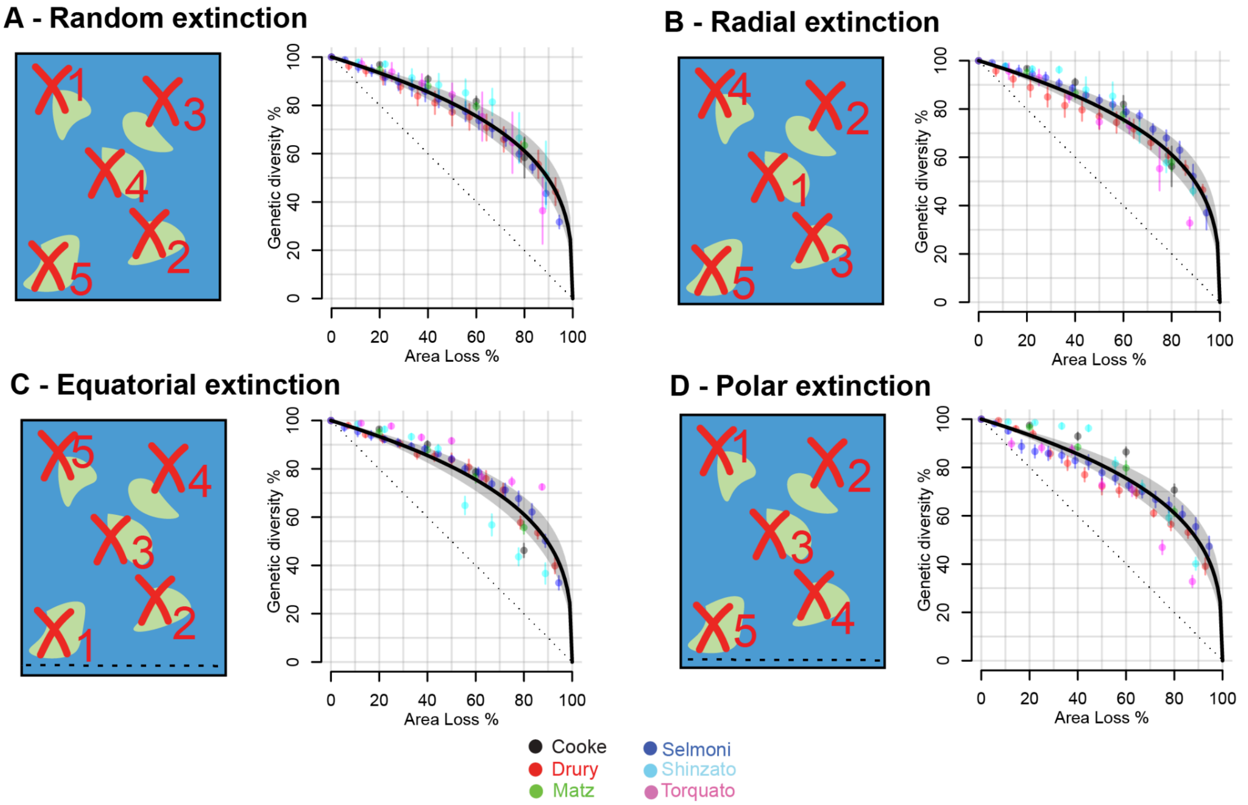


**Figure S21. Genetic extinction by area loss simulations.**

Shown are the results of stochastic extinction simulations across the *Acropora* datasets (colors in the graphs). In each simulation scenario, the study area of the datasets was progressively reduced, removing individuals from the datasets and counting the amount of mutations (genetic diversity %) lost. The different scenarios show distinct spatial patterns of area loss (red crosses in the drawings display the order of extinctions): (A) reefs are lost following a random order, (B) extinction starts at a given reef, then expands to the neighboring ones; (C) extinction moves from low to high latitudes; (D) extinction moves from high to low latitudes. The black line in the graph represents the expected loss of genetic diversity by area loss according to the average *Acropora* z-mar (with interval of confidence in the shaded area).

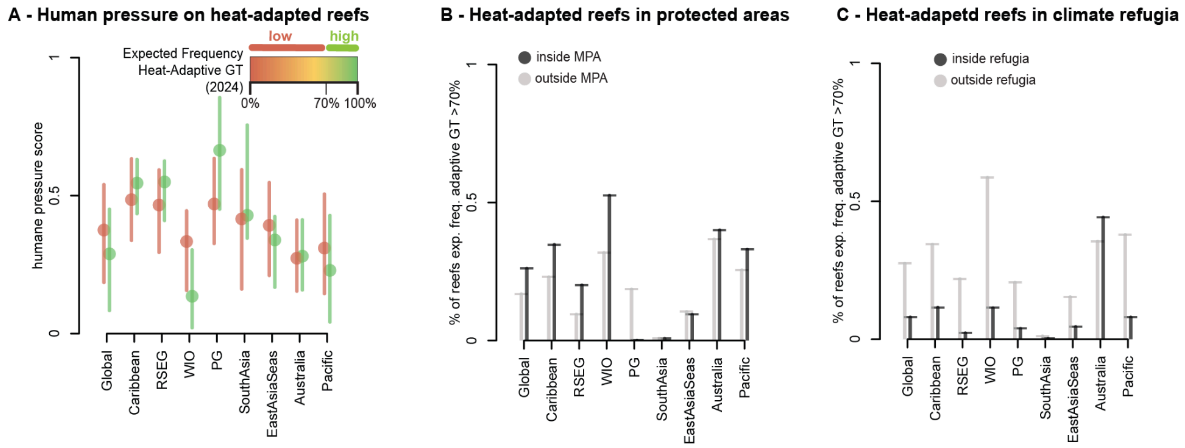


**Figure S22. Heat-adapted reefs in conservation and management strategies in 2020.**

Shown are three metrics of the conservation status of heat-adapted reefs (*i.e.,* reefs with an expected frequency of heat-adaptive genotypes ≥70%) and non heat-adapted reefs (*i.e.*, expected frequency <70%) across eight oceanic regions. (A) shows the human pressure index (0: low pressure, 1: high pressure) in heat-adapted vs. non heat-adapted reefs. (B) shows the fraction of heat-adapted and non-heat adapted reefs that are inside and outside Marine Protected Areas (MPA). (C) shows the fraction of heat-adapted and non heat-adapted reefs that are included or excluded in a portfolio of 50 climate refugia - *i.e.*, reef areas that are projected to withstand climate change, based on a previous study on connectivity and historical and forecasted climate (*53*).
